# Molecular mode of action of an Acyl Protein thioesterase

**DOI:** 10.1101/2020.06.18.157545

**Authors:** Laurence Abrami, Martina Audagnotto, Sylvia Ho, Maria Jose Marcaida, Francisco S. Mesquita, Muhammad U. Anwar, Patrick A. Sandoz, Giulia Fonti, Florence Pojer, Matteo Dal Peraro, F. Gisou van der Goot

## Abstract

Many biochemical reactions occur at the membrane interfaces. The proper control of these reactions requires spatially and temporally controlled recruitment of protein complexes. These assemblies are largely regulated by post-translational modifications and a frequent one is S-acylation, which consists of the addition of medium length acyl chains. Reversibility of this modification is ensured by acyl protein thioesterases (APTs), which are poorly understood enzymes. Using a combination of computational, structural, biochemical, and cellular approaches, we dissect the mode of action of a major cellular thioesterase, APT2 (LYPLA2). We show that for APT2 to encounter its targets, it must interact with membranes by two consecutive steps, the insertion of a hydrophobic loop and subsequent S-acylation by the ZDHHC3 or ZDHHC7 palmitoyltransferases. Once bound, APT2 deforms the lipid bilayer to extract the acyl chain bound to its substrate, capturing it in a hydrophobic pocket and allowing hydrolysis. Deacylation releases APT2, allowing it to bind to other membranes, but also renders it vulnerable to ubiquitination and proteasomal degradation. This molecular understanding of APT2 paves the way to understand the dynamics of APT2-mediated depalmitoylation throughout the endomembrane system.

## INTRODUCTION

Eukaryotic cells are complex factories, with millions of separate reactions occurring simultaneously to control every aspect of how that cell functions and behaves with respect to other cells. This requires the exquisite spatial and temporal control of proteins, often through post-translational modifications. One of the most frequent modifications, affecting 10 to 20% of the human proteome, is the S-acylation (Khoury *et al*, 2011; Blanc *et al*, 2019), which can modify a plethora of signaling molecules, such as the EGF receptor or Ras, adhesion molecules, and many transporters. S-acylation is also a key modification in the life cycle of viruses (Gadalla & Veit, 2020) and parasites (Brown *et al*, 2017). This lipidation may affect the trafficking, function, or turnover rate of proteins, and therefore its reversibility is an essential part of its regulatory capacity (Zaballa & van der Goot, 2018). Little is known, however, about acyl protein thioesterases (APTs), the proteins responsible for removing the acyl chains. Therefore, we herein focused on understanding the mode of action of APT2, a major cytosolic thioesterase, that is involved in the palmitoylation cycle of a variety of proteins, such as the TNF receptor (Zingler *et al*, 2019), the melanocortin 1 receptor (Chen *et al*, 2019), and the palmitoytransferase ZDHHC6 (Abrami *et al*, 2017).

Similar to other lipid modifications, S-acylation modifies the lipophilicity of proteins, thereby affecting their ability to interact with membranes or membrane domains. S-acylation is unique amongst lipid modifications in its reversibility (Conibear & Davis, 2010; Fukata *et al*, 2016; Lemonidis *et al*, 2015; Salaun *et al*, 2010). During enzymatic S-acylation, medium chain fatty acids are added to cysteine residues through the action of acyltransferases of the ZDHHC family, which are transmembrane proteins with an active site on the cytosolic side of the membrane. Regulatory removal of the fatty acid is mediated by cytosolic APTs (Lin & Conibear, 2015; Won *et al*, 2018), wherein all identified members belong to the α/β hydrolase super family. APT2 is a soluble globular enzyme that can undergo S-acylation at its N-terminus on Cys2 (Kong *et al*, 2013; Vartak *et al*, 2014). This is thought to bring APT2 to the membranes that contain its substrates for encounter with and subsequent deacylation of that substrate. Even though the structure of an inhibitor-bound form of APT2 has been solved (Won *et al*, 2016), how APT2 interacts with the membrane and how it catalyzes the release of the bound fatty acid is not known. Because it has been shown that single acylation of a protein is insufficient to allow stable association with membranes (Shahinian & Silvius, 1995), we anticipate that APT2 must have an additional mode of membrane interaction, which could be dimerization, a second lipid modification, or a lipid-binding motif within the protein.

To investigate how APT2 binds to membranes and brings about the deacylation of its targets, we first choose a structural approach combining X-ray crystallography and molecular dynamics (MD) simulations. Using this, we identified a loop, coined the ß tongue, which mediates membrane interaction both *in vitro* and *in vivo* in a step that precedes and is necessary for S-acylation of APT2 to occur. We identified the ZDHHC palmitoyl transferases involved and show that the sequential membrane association process controls both the turnover rate of APT2 and its activity. Next, we identified a hydrophobic pocket within APT2 that captures the acyl chain prior to hydrolysis. Molecular dynamics simulations further indicate that APT2 can deform the lipid leaflet to which it binds, to extract the substrate-bound acyl chain from the membrane, position it in a hydrophobic pocket allowing the thioester bond to be cleaved in the catalytic site. Finally, we show that cells tightly control the concentration of soluble APT2, through a mechanism built within the ß tongue, where a sensitive ubiquitination site is protected in the membrane bound state. Altogether, these results on APT2 provide detailed and previously unknown information about how this thioesterase mediates de-acylation and is regulated in the cell, helping broaden our picture of how S-acylation cycles control cellular processes.

## RESULTS

### APTs are monomers in solution

Our first goal was to perform a crystallization study on full length APTs, as opposed to previous N-terminal truncated versions (Devedjiev *et al*, 2000) and to compare WT and mutants. APT1 and APT2 are 68% identical and were found to have essentially the same structure (Wepy *et al*, 2019; Won *et al*, 2016). As previously reported (Won *et al*, 2016), however, APT1 is more stable, and was more amenable to crystallization. Therefore, we performed our crystallization studies on APT1. We produced full-length, human, wildtype (WT) APT1, as well as palmitoylation deficient (C2S) and catalytically inactive (APT1-S119A and APT2-S122A) mutants in *E. coli*. All variants could be successfully crystalized (**Supplementary Table 1**). APT1 has a near-canonical α/β hydrolase fold characterized by the presence of the typical catalytic triad (i.e., Ser119, Asp174, and His208, **Fig. 1ab**) and a central β-sheet (labeled β2 to β8 in **Fig. 1ab**), which is connected to five α-helices (labeled A to F) and four smaller α-helices (labeled G1 to G4) (Devedjiev *et al*, 2000). Importantly, when compared to the canonical α/β hydrolase fold, both APT1 and APT2 have an atypical insertion of four short antiparallel β strands (labeled S1-S4, **Fig. 1b**) organized into loop-like structures that, as described below, are critical to their protein acylthioesterase function.

**Figure 1:**
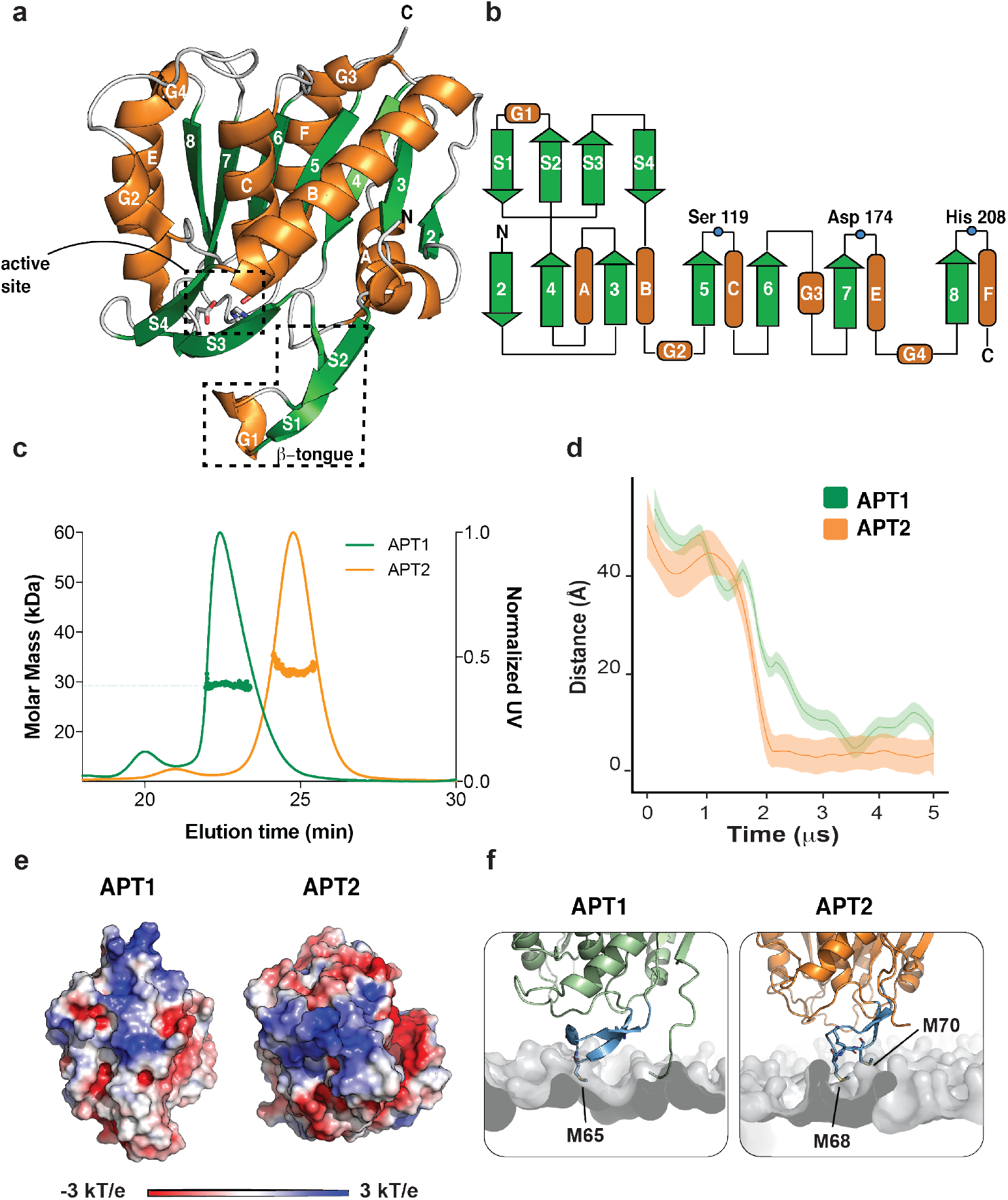
Structure and membrane interaction of APT2. **a.** Ribbon diagram of the secondary structure elements. Seven β strands (green arrows) are numbered sequentially from 2 to 8. APT2 contains an insertion atypical of the α/ß hydrolase fold corresponding to four short β strands named S1-S4. Five α helices (orange) are labeled from A to F while four short α helices are labeled from G1 to G4. The catalytic active site and the β-tongue are contoured in black. **b.** Topology of the APT enzyme. **c.** SEC-MALS analysis shows that both APT1 and APT2 elute as single peaks that correspond to the monomeric forms, with molar masses of 29 and 32 kDa, respectively. **d.** Distance plot that monitors the APT1 (green) and APT2 (orange) membrane interaction over the MD simulation time. **e.** Electrostatic surface potential of APT1 (left) and APT2 (right) shows the highly positive surface (blue) that promotes the interaction with the lipid bilayer. **f.** Close-up view of the enzyme-protein interaction for APT1 (left) and APT2 (right). The β-tongue is highlighted in blue. The side chains of the residues involved in the membrane interaction are shown as blue sticks.

The APT1 variants crystallized into different forms, with space group P4_1_2_1_2 for WT, P2_1_2_1_2 for the C2S mutant, and P6_4_ for the S119A mutant (**Supplementary Table 1**). The average RMSD for the 230 common Cα atoms between our structures and the first published APT1 structure (PDB id: 1FJ2, Devedjiev *et al*, 2000) was ~1.2 Å. The asymmetric units of the crystals contained 2 or 6 monomers (**Supplementary Table 1**). For each structure, EPPIC (Duarte *et al*, 2012) predicted the possible existence of dimers, as was previously reported. Our WT and C2S X-ray structures showed a similar intermolecular interface (RMSD ~1.1 Å). In contrast, the previously solved APT1 structure (PDB id: 1FJ2, Devedjiev *et al*, 2000) and the catalytically inactive APT1 mutant (S119A) had a largely different conformation (RMSD ~14.2 Å) (**Supplementary Fig.1**). Principal component analysis based on atomistic MD simulations of the WT APT1 structure highlighted a large rotational motion centered at the supposed dimeric interface (**Supplementary Fig.1**). Crystallographic evidence and MD simulations thus indicate that the dimeric arrangement of APT1 is intrinsically flexible. Interestingly, these dimerizing interfaces also included the APT1 catalytic pocket such that it was partially occluded by the adjacent protomer. Similar observations where made regarding APT2. Backbone RMSD calculations on superimposed APT2 structures (PDB: 6BJE and 5SYN; RMSD 14Å, **Supplementary Fig.1cd**) combined with structural comparison and principal component analysis indicate a flexible dimer interface. These observations indicate that both APT1 and APT2 have dimeric conformations that are probably transient, suggestive of a monomeric active form of the enzyme.

To directly address the stoichiometry of APTs in solution, we measured molecular weights using size-exclusion chromatography (SEC) coupled to multi-angle light scattering (MALS). The molecular mass determined by SEC-MALS was 29 kDa for APT1 and 32 kDa for APT2 (**Fig. 1c**), close to the theoretical molecular weights of monomers (25 and 27 kDa, respectively), indicating that the enzymes are monomeric in solution at physiological concentrations. These observations reconcile the finding of a flexible, heterogeneous dimer interface in APT1 and APT2 under crystallization conditions and suggest that the APTs act as monomeric entities in solution and most likely also when interacting with membrane surfaces, as the catalytic site would be otherwise occluded.

### APTs are predicted to have intrinsic membrane binding affinity

The target proteins of APTs are S-acylated proteins, so to reach their membrane-associated substrates, APTs must be near membranes. We explored whether APTs can interact with membranes directly using coarse-grained MD (CG-MD) simulations in the presence of a palmitoyl-oleoyl-phosphatidylcholine (POPC) bilayer model. The simulations showed that APT1 and APT2 invariably interact with the membrane bilayer (**Fig. 1d**) and always via the same surface (**Fig. 1ef**), which corresponds to the intermolecular interface observed in the APT X-ray structures (**Supplementary Fig.1**). Electrostatic calculations showed that the membrane-interacting surfaces of APT1 and APT2 contained highly positively charged regions that likely promote the initial association with the membrane by long-range electrostatic attraction (**Fig. 1e**). APT1 has two distinct positively charged regions (region A: Arg13, Lys14, His43 and Lys45; region B: His23, His30 and His50) (**Fig. 1e,** left), while APT2 has a single larger area (Arg59, His55, His28, His170, His208) (**Fig. 1e,** right), allowing both proteins to regularly interact with the membrane on a μs timescale. Interestingly, once close to the membrane, APT1 and APT2 use the S1 and S2 strands and the G1 helix, which are mostly hydrophobic, to anchor the enzyme to the membrane (**Fig. 1f**). S1, S2 and G1 are part of the atypical insertion that APTs have when compared to classical α/β hydrolases (**Fig. 1b**). By analogy to a similar membrane-anchoring loop in the pore-forming toxin cytolysin A (ClyA) (Mueller *et al*, 2009), we coined the S1-G1-S2 loop the ß tongue.

### *In vitro* APT membrane binding is mediated by the ß tongue

To test the prediction that APTs have intrinsic membrane binding activity, we incubated purified APT1 and APT2 with liposomes composed of phosphatidylcholine: phosphatidylserine: phosphatidylethanolamine (PC:PS:PE 2:2:1). The mixtures were subsequently submitted to a sucrose flotation step gradient to separate vesicle-bound from unbound protein. While the proteins remained in the bottom fraction in the absence of liposomes, they were recovered at the top interface when liposomes were present (**Fig. 2a**), indicating an interaction with the membrane. Analysis of the APT-bound liposomes by electron microscopy confirmed the presence of a layer of proteins decorating the surface of the liposomes (**Fig. 2b**).

**Figure 2:**
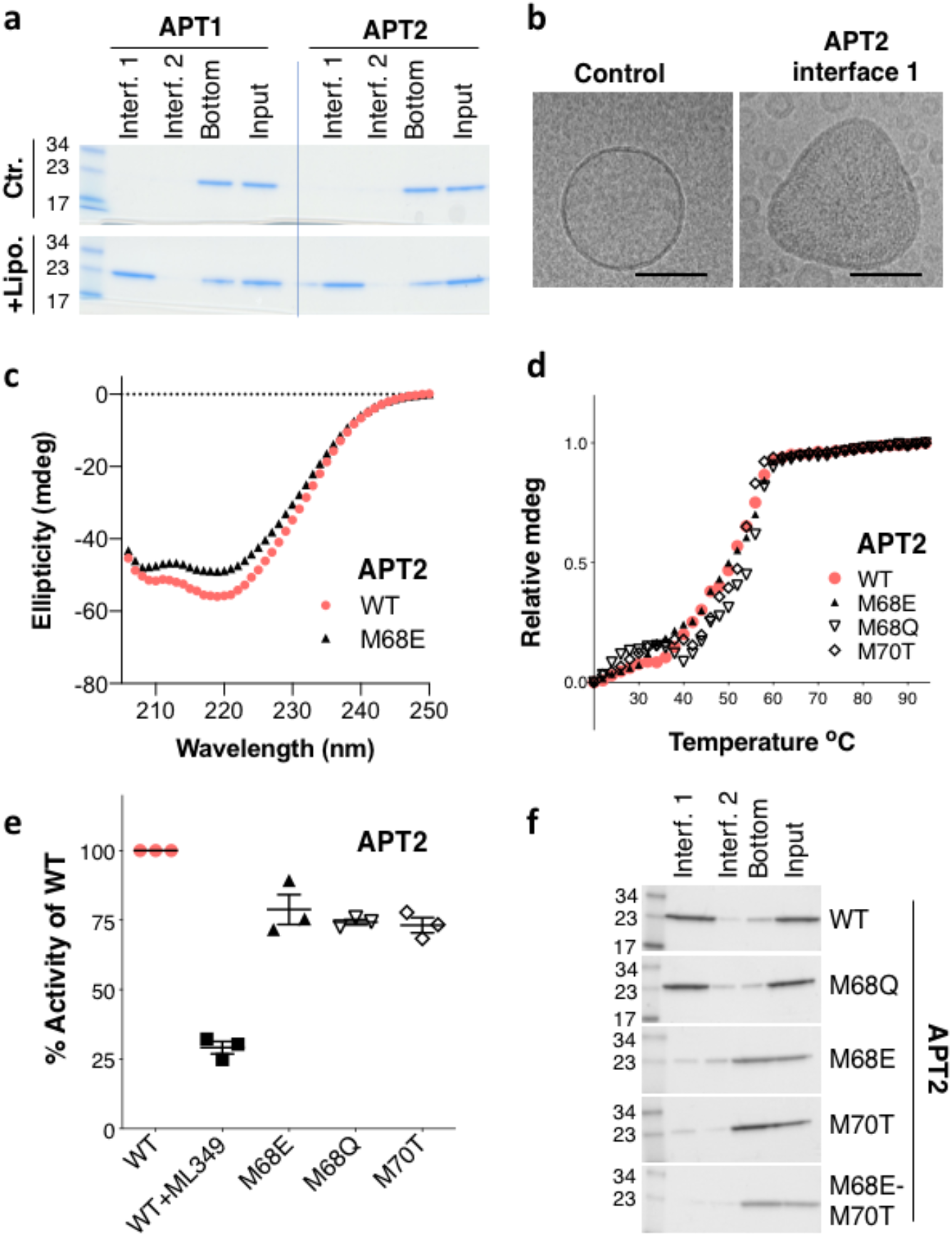
ß–tongue-mediated membrane binding of APTs. **a.** Purified WT APT1 and WT APT2 protein were incubated with liposomes and applied to the bottom of a step sucrose gradient. The interfaces were collected from top to bottom, loaded on an SDS-PAGE gel, and stained with Coomassie blue. **b.** Interface 1 (Interf. 1) for WT–APT2-loaded liposomes was analyzed by negative staining and cryo EM. Bar = 100 μm. **c.** Circular dichroism spectra of WT APT2 and the ß tongue M68E mutant. **d.** Thermal denaturation profiles of WT APT2 or the ß tongue mutants as monitored by circular dichroism at 222 nm. The normalized ellipticity at 222 nm is plotted against temperature. **e.** The thioesterase activity of WT APT2 or the ß tongue mutants was determined at 60 min after the addition of substrate and detergent. APT2-specific inhibitor ML349 was included as a positive control. Technical replicates were averaged within each experiment, and then each experiment was normalized to WT. The average of each experiment was graphed. Two-tailed two-sample unequal variance t-tests were performed on the raw values normalized to the plate. **f.** WT and different APT2 mutants were incubated with liposomes and applied to the bottom of a sucrose gradient. The different interfaces from the top to bottom were collected and loaded on an SDS-PAGE and stained with Coomassie blue.

To analyze the importance of the ß tongue in this interaction, we generated various mutants. We focused mostly on APT2 because, as opposed to APT1 that is most abundant in mitochondria (Kathayat *et al*, 2018), APT2 is found on the cytosolic side of membranes and is thus topologically able to revert the action of ZDHHC enzymes. We produced the M68Q, M68E, M70T, and M68E-M70T APT2 mutants as well as the M65E APT1 mutant. Secondary structure analysis of the APT2 mutants using circular dichroism (CD) in the far UV range indicated that the ß tongue mutations did not significantly affect the APT2 secondary structure (**Fig. 2c**). We also performed unfolding studies measuring the change in ellipticity at 222 nm as a function of temperature. The melting temperature was similar for WT and mutant APT2s (**Fig. 2d**), indicating that the stability of the proteins was preserved. Using a colorimetric assay, we verified that the *in vitro* deacylating activity of APT2 was not drastically affected by the mutations (**Fig. 2e**). Finally, we tested their liposome binding capacity. While the M68Q behaved like WT APT2, all other mutants failed to stably bind to liposomes (**Fig. 2f**). Similarly, the M65E APT1 ß tongue mutation impaired liposome binding (**Supplementary Fig.1e**). Altogether, these experiments confirm the predictions from the MD simulations that APTs have intrinsic membrane binding affinity driven by the ß tongue.

### Membrane-anchoring-deficient APT2 mutants are targeted to the proteasome

We next analyzed the APT2 ß tongue mutants upon expression in cells. To our surprise, those that failed to bind liposomes were barely expressed upon transient transfection in HeLa cells (**Fig. 3a**, control). Expression could be rescued by the proteasome inhibitor MG132 (**Fig. 3b**, +MG132), suggesting that the mutants were synthesized by the cell but rapidly degraded. To confirm this, we performed ^35^S-Cys/Met metabolic pulse-chase experiments. While the apparent half-life of WT APT2 after a 20 min pulse was approximately 5 h, it dropped to <2 h for the M68E and M70T mutants (**Fig. 3b**).

**Figure 3:**
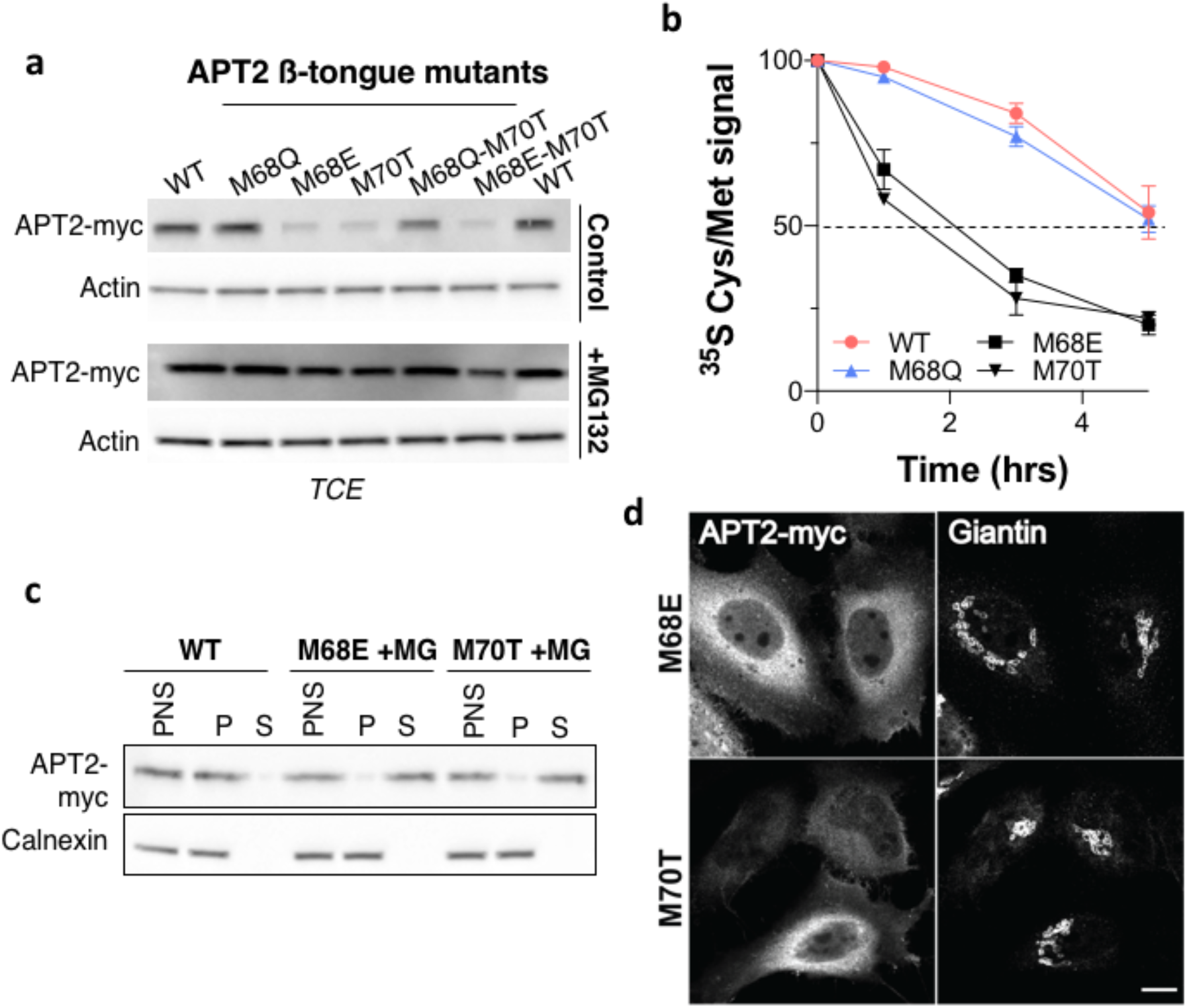
APT2 ß tongue mutants are cytosolic and unstable. **a.** Cells expressing WT or mutant APT2-myc for 24 h were incubated for 4 h with MG132. Protein extracts (40 μg) were separated via SDS-PAGE and analyzed by immunoblotting with anti-myc antibodies. Anti-actin antibodies were used as a loading control. **b.** HeLa cells were transfected with different APT2-myc constructs for 24 h. Cells were pulsed with ^35^S Cys/Met for 20 min and were chased for the indicated time before immunoprecipitation and SDS-PAGE. Degradation kinetics were analyzed by autoradiography and were quantified using the Typhoon Imager. ^35^S-Met/Cys incorporation was quantified for each time point and was normalized to protein expression levels. ^35^S-Met/Cys incorporation was set to 100% for t = 0 after the 20-min pulse, and all chase times were expressed relative to this (n = 3, error bars represent standard deviation). **c.** HeLa cells were transfected with plasmids encoding WT or mutant APT2-myc for 24 h. Where applicable, cells were incubated for 4 h with MG132. Post-nuclear supernatants (PNS) were prepared and ultra-centrifuged to separate membrane (P) from cytosolic (S) fractions. Equal volumes were loaded on a 4–20% gradient SDS-PAGE gel and were analyzed by immunoblotting with anti-myc and anti-Calnexin antibodies. **d.** Confocal microscopy images of HeLa cells transfected with plasmids encoding APT2-myc mutants for 24 h. Cells were incubated for 4 h with MG132 and were immunolabeled for APT2-myc and giantin. Scale bar: 10 μm.

To test the membrane binding capacity of the mutants, we expressed APT2 variants and treated cells with MG132 for 4 h, before submitting post-nuclear supernatants (PNS) to high speed centrifugation. Analysis of the pellets and the supernatants showed that while the WT protein was mostly found in the pellet, the M68E and M70T mutants were exclusively found in the supernatant (**Fig. 3c**). These observations were confirmed by immunofluorescence analysis, which showed a cytosolic-like staining of the mutants (**Fig. 3d**). This indicates that ß tongue APT2 mutants, which failed to bind to liposomes, also remained cytosolic when expressed in cells and underwent rapid proteasome-mediated degradation.

### Stable membrane association of APT2 requires S-acylation of Cys2

The cytosolic distribution of APT2 ß tongue mutants differs significantly from that of the WT protein, which shows a marked accumulation on the Golgi apparatus (**Fig. 4a**). This Golgi localization was previously reported and found to be dependent on Cys2 (Kong *et al*, 2013; Vartak *et al*, 2014) and its likely S-acylation. Using the C2S mutant, we confirmed that palmitoylation occurs on Cys2 (**Supplementary Fig. 2a**) and that the WT protein accumulates on the Golgi in a Cys2-dependent manner (**Fig. 4a**). In agreement with the microscopy (**Fig. 4a**), the C2S mutant was mostly found in the cytosolic fraction after high speed centrifugation of the PNS, in contrast to the WT protein that was mostly found in the pellet, consistent with a membrane association (**Fig. 4b**).

**Figure 4:**
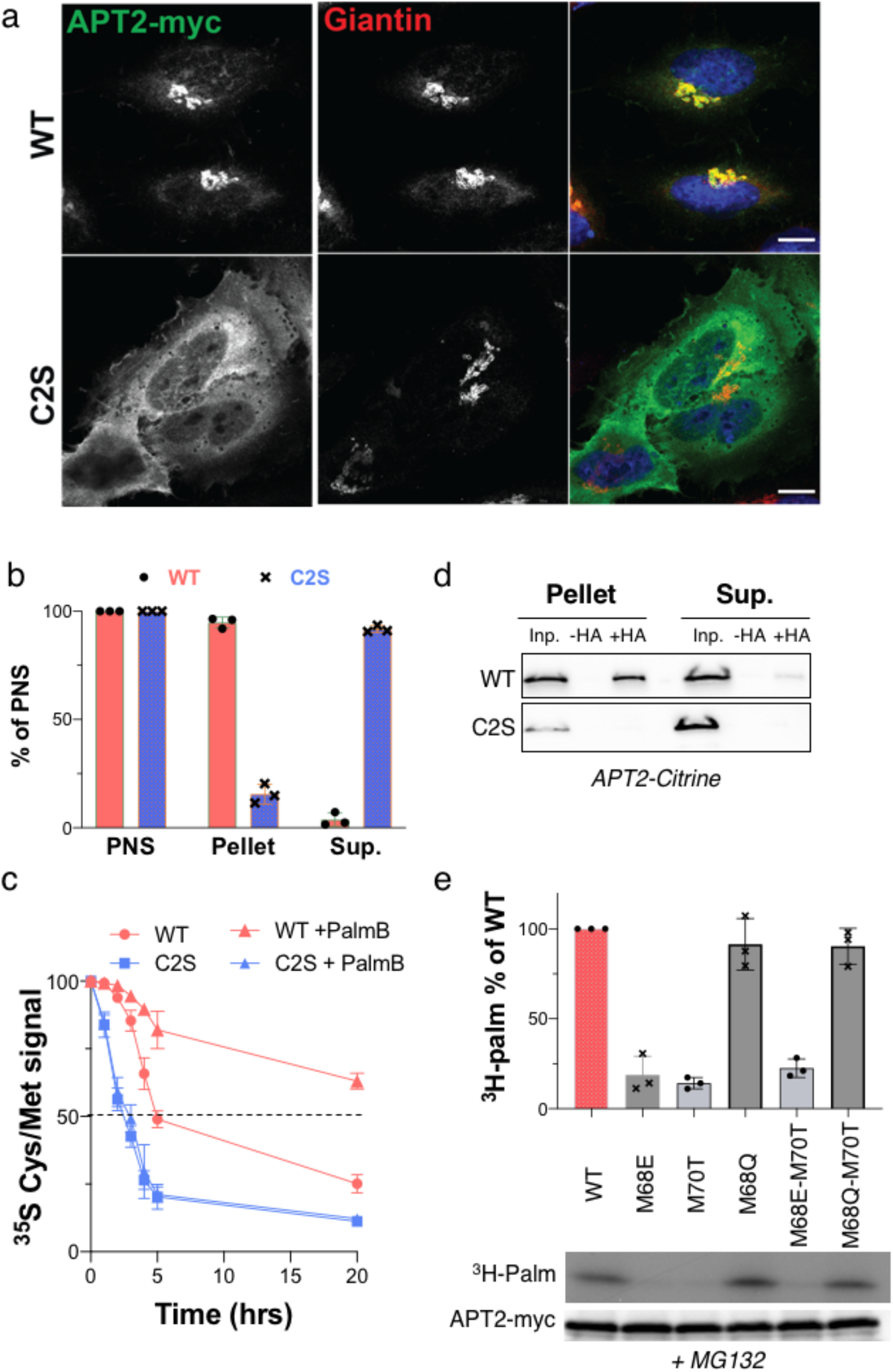
Effect of S-acylation on APT2 localization and stability. **a.** Cellular localization of APT2 mutants. Confocal microscopy images of HeLa cells transfected with plasmids encoding WT or C2S APT2-myc for 24 h and immunolabeled for APT2 and giantin. The nuclei were stained with Hoechst. Scale bar: 10 μm. **b.** HeLa cells were transfected with plasmids encoding WT or C2S APT2 for 24 h. PNS were ultra-centrifuged to separate the membrane (Pellet) and cytosolic (Sup.) fractions. Equal volumes were analyzed by SDS-PAGE. Quantification of the APT2 membrane association was performed by setting the amount of APT2 in PNS to 100%, and the amount of WT or C2S APT2 in the pellet or supernatant were expressed relative to this (n = 3, error bars represent standard deviation). **c.** HeLa cells were transfected with different APT2-myc constructs for 24 h, treated or not for 4 h with Palmostatin B, pulsed with ^35^S Cys/Met for 20 min and then chased for the indicated time before immunoprecipitation and SDS-PAGE. Degradation kinetics were analyzed by autoradiography and were quantified using the Typhoon Imager. ^35^S-Met/Cys incorporation was quantified for each time point and was normalized to protein expression levels. ^35^S-Met/Cys incorporation was set to 100% for t = 0 after the 20-min pulse, and the different chase times were expressed relative to this (n = 3, error bars represent standard deviation). **d.** HeLa cells were transfected with plasmids encoding WT or C2S APT2-cintrine constructs for 24 h. PNS were ultra-centrifuged to separate the membrane (Pellet) and cytosolic (Sup.) fractions. The amount of palmitoylated protein was determined using Acyl-RAC. For each input (Inp.), palmitoylated proteins were detected after hydroxylamine treatment (+HA). Equal volumes were by analyzed SDS-PAGE and immunoblotting with anti-GFP antibodies. **e.** HeLa cells were transfected with plasmids encoding WT or the indicated APT2-myc mutants for 24 h. Cells were treated for 4 h with MG132 and were then metabolically labeled for 3 h at 37°C with ^3^H-palmitic acid. The proteins were extracted, immunoprecipitated with myc antibodies, separated via SDS-PAGE, and analyzed by autoradiography (^3^H-palm), which was quantified using the Typhoon Imager or by immunoblotting with anti-myc antibodies. The calculated value of ^3^H-palmitic acid incorporation into WT APT2 was set to 100%, and mutants were expressed relative to this (n = 3, error bars represent standard deviation).

Since ß tongue mutants are rapidly degraded in cells, we wondered whether S-acylation, which also affects membrane binding, would similarly influence APT2 turnover rates. We performed ^35^S-Cys/Met metabolic pulse-chase experiments on WT and C2S in the presence or absence of the general protein deacylation inhibitor Palmostatin B. The C2S mutant had a shorter apparent half-life (≈3 h) than the WT protein (≈5 h) (**Fig. 4c** and **S3b**). As expected, the decay of C2S was not affected by Palmostatin B treatment. In contrast, degradation of the WT protein was drastically slowed by Palmostatin B, with the half-life increasing to more than 20 h (**Fig. 4c** and **S2bc**). This is not due to an effect of membrane binding on protein stability *per se*, since the melting curve of APT2 was the same in solution and when bound to liposomes (**Supplementary Fig.2d**). Instead, the cellular instability likely indicates that the acyl modification affects recognition of APT2 by a cytosolic quality control mechanism.

The S-acylation of APT2 is thus required for stable membrane association, and this modification drastically affects APT2 turnover rates in the cellular context. The strongly stabilizing effect of Palmostatin B also suggests that APT2 undergoes significant depalmitoylation in cells, an issue we will address in future studies.

### APT2 membrane association occurs through a two-step mechanism

We next investigated the steady state S-acylation status of APT2 in cells by performing a capture assay for S-acylated proteins termed Acyl-RAC. We analyzed both the cytosolic and pellet fractions of the PNS. Acylated WT APT2 was readily detectable in the pellet fraction, but not in the supernatant (**Fig. 4d**), indicating that the fraction of soluble APT2 is not S-acylated.

Given that the ß tongue mutants were also found in the soluble fraction, this suggests that despite the presence of the cysteine at position 2, these mutants might not be able to undergo S-acylation. To test this, ß tongue mutants were expressed in HeLa cells and their degradation was prevented by MG132 treatment. The incorporation of ^3^H-palmitate was monitored. While palmitoylation of WT APT2 was readily observed, the membrane-binding-deficient ß tongue mutants did not undergo detectable palmitoylation (**Fig. 4e**).

These observations altogether indicate that APT2 undergoes a two-step membrane-binding process in the cell: (i) electrostatic interactions bring APT2 to the membrane, where the ß tongue dips into the membrane to hold the APT2 temporarily in place. This is necessary for APT2 to encounter its S-acylating enzyme(s), (ii) leading to the lipidation of Cys2 and a stable membrane association. The effect of palmostatin B on APT2 stability further indicate that APT2 can undergo deacylation and membrane release.

### APT2 S-acylation is required for in vivo activity and is mediated by ZDHHC3 and 7

We next analyzed whether S-acylation is important for APT2 activity, as APT2 is a soluble enzyme and its targets are membrane bound. As a readout for activity, we monitored the APT2-dependent depalmitoylation of one of its targets, the zDHHC6 protein. Endogenous APT2 expression in cells was silenced and subsequently complemented with either WT or C2S APT2. The cells were pulsed for 2 h with ^3^H-palmitate and chased for 0 or 3 h to allow APT2-mediated depalmitoylation. zDHHC6 was then immunoprecipitated, and its palmitoylation level was monitored by autoradiography. Upon expression of WT APT2, depalmitoylation of ZDHHC6 was observed at 3 h (**Fig. 5a**), as expected for the normal function of APT2. However, C2S APT2 failed to mediated ZDHHC6 depalmitoylation (**Fig. 5a**), indicating that the S-acylation site at position 2 is necessary for APT2 activity.

**Figure 5:**
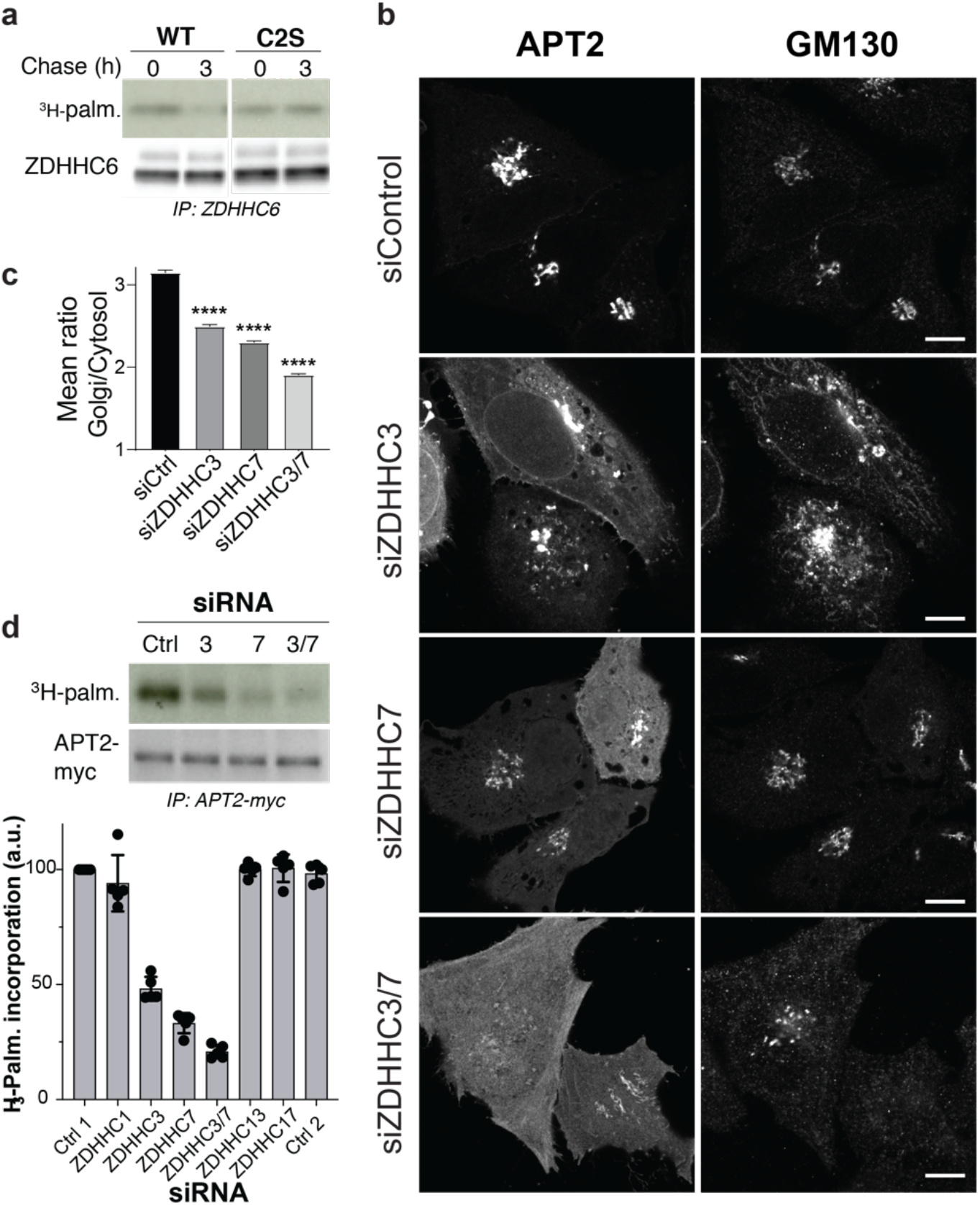
APT2 S-acylation is mediated by ZDHHCs 3 and 7 and is required for activity. **a.** HeLa cells were silenced for 3 days with APT2 RNAi (in 3’ non-coding sequence) transfected with plasmids encoding myc-tagged WT ZDHHC6 and APT2 WT or C2S for 24 h. The cells were then metabolically labeled for 2 h at 37°C with ^3^H-palmitic acid and were chased for 3 h in new complete medium. Proteins were extracted, immunoprecipitated with anti-myc antibodies, subjected to SDS-PAGE, and analyzed by autoradiography (^3^H-palm) or by immunoblotting with anti-myc antibodies. **b.** Confocal microscopy images of HeLa cells transfected with siRNAs against ZDHHC3, 7 or both, plasmids encoding APT2-citrine for 24h, and immunolabelled for GM130. Scale bar: 10 μm **c.** High-throughput automated immunofluorescence quantification of HeLa cells processed as in **b**, depicting the ratio of APT2 mean intensity values between the Golgi (marked by GM130) and the cytosol (cell mask without nuclei and Golgi). At least 2200 cells were analyzed per condition and results are mean ± SEM p****<0.0001. Equivalent results were obtained for two independent analysis. **d**. HeLa cells were transfected with siRNAs against ZDHHC3, 7 or both and a plasmid encoding APT2-myc, then metabolically labeled for 2 h at 37°C with ^3^H-palmitic acid. Proteins were extracted, immunoprecipitated with anti-myc antibodies, subjected to SDS-PAGE, and analyzed by autoradiography (^3^H-palm) or by immunoblotting with anti-myc antibodies (n = 5, error bars represent standard deviation).

Given the importance of S-acylation on APT2 localization, cellular stability, and activity, we performed a screen to identify the ZDHHC enzyme(s) responsible for modifying APT2. Because APT2 accumulates on the Golgi when acylated, we could screen by fluorescence microscopy. We first silenced ZDHHC enzymes by dividing them into 6 pools. Silencing pool M1, containing siRNAs against ZDHHC 1, 3, 7, 13, and 17, led to the Golgi release of APT2 (**Supplementary Fig.3**). We subsequently silenced each ZDHHC enzyme individually and found that none of them led to the complete release of APT2 from the Golgi. We therefore silenced them in pairs and found that the simultaneous silencing of ZDHHC3 and 7 (**Fig. 5bc** and **S3**), but not for example 13 and 17 (**Supplementary Fig.3**), led to APT2 release. We could confirm ZDHHC3- and 7-dependent palmitoylation of APT2 by ^3^H-palmitate incorporation (**Fig. 5d**).

### Control of APT2 turnover by the ß tongue

The dual interaction of APT2 with the membrane, via the ß tongue and through lipidation of Cys2, allows stable membrane association, though also appears to protect APT2 from proteasomal degradation. To understand this protective mechanism, we searched for lysine residues that could be targets for ubiquitination. APT2 contains 6 lysines in total, one of which appears ideally positioned at the very tip of the ß tongue, Lys-69 (**Fig. 6a**). We generated K69R mutants for the WT protein and the ß tongue mutants. Remarkably, the K69R mutation led to a full rescue of the expression of the membrane-binding-deficient ß tongue mutants and a loss of the ubiquitination signal (**Fig. 6b**). Introducing the K69R mutation also led to an increase in the expression of the C2S mutant (**Fig. 6b**). These observations reveal that APT2 has a built-in mechanism whereby soluble APT2, through the exposure of Lys-69, can be ubiquitinated and targeted to degradation, while membrane-bound APT2 dips Lys69 in the membrane, hiding if from the ubiquitin ligase.

**Figure 6:**
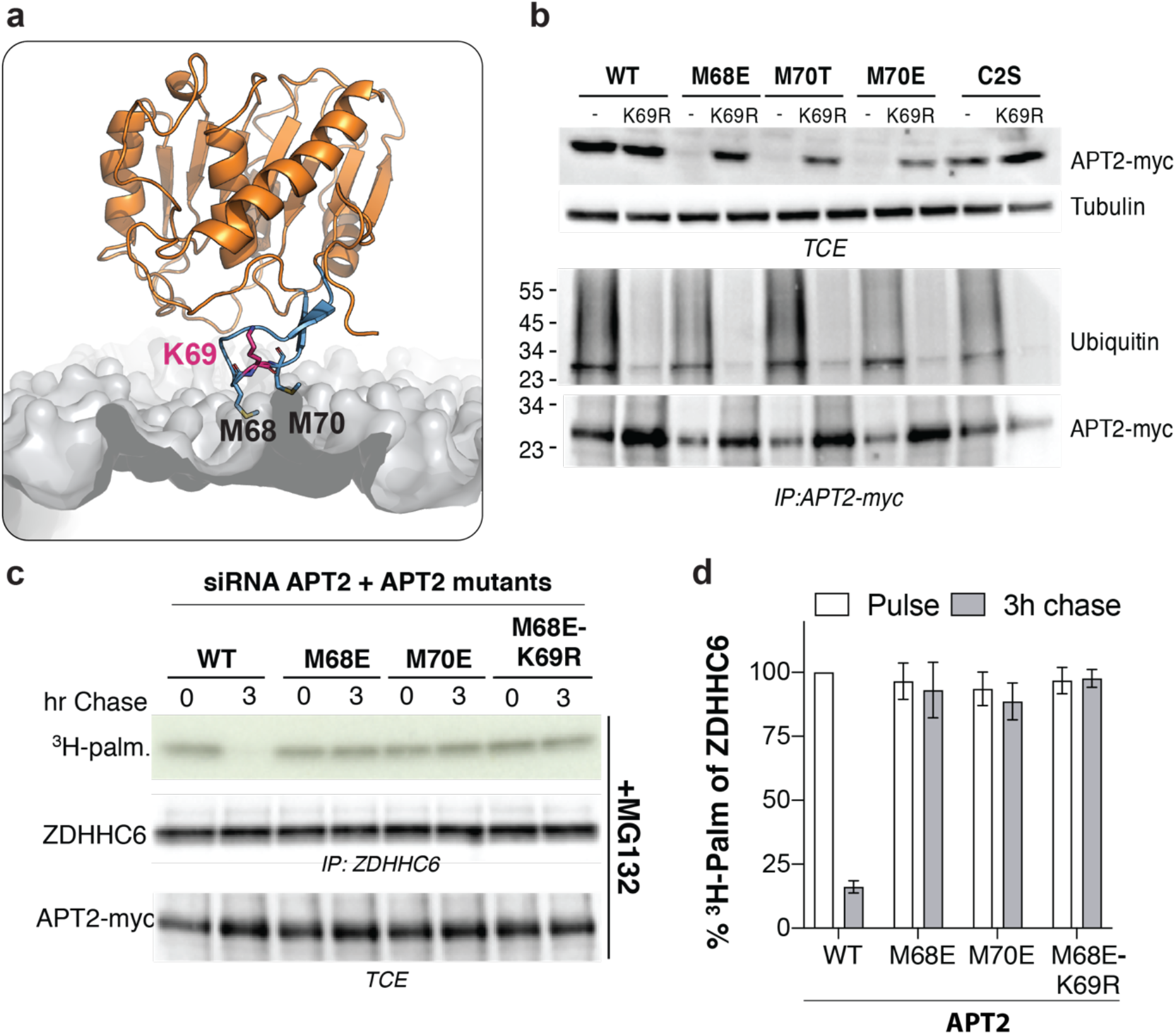
APT2 ß tongue mutants undergo ubiquitination on Lys-69 but not S-acylation. **a**. Ribbon diagram of membrane-bound APT2. **b.** HeLa cells expressing WT or mutant APT2 for 24 h were lysed and immunoprecipitated with anti-myc agarose beads overnight. Protein extracts (40 μg) and immunoprecipitated products were separated via SDS-PAGE and analyzed by immunoblotting with anti-myc or anti-ubiquitin antibodies. Anti-tubulin antibodies were used as a loading control. **c.** HeLa cells were silenced for 3 days with APT2 siRNA (in 3’ non-coding sequence) and were transfected with plasmids encoding myc-tagged WT ZDHHC6 and the indicated APT2 constructs for 24 h. Cells were incubated for 4 h with MG132 and then were metabolically labeled for 2 h at 37°C with ^3^H-palmitic acid and chased for 3 h in new complete medium. Proteins were extracted, immunoprecipitated with anti-myc antibodies, subjected to SDS-PAGE, immunoblotted with anti-myc antibodies, and analyzed by autoradiography (^3^H-palm). The total extracts (40 μg) were immunoblotted with anti-myc antibodies to detect WT and mutant APT2. **d.** Quantification of ^3^H-palmitic acid incorporation into ZDHHC6 in the presence of different APT2 mutants. Values were normalized to protein expression level. The calculated value of ^3^H-palmitic acid incorporation into ZDHHC6 with WT APT2 was set to 100%, and the mutants were expressed relative to this (n = 3, error bars represent standard deviation).

Rescuing the expression of the ß tongue mutants by the K69R mutation was however insufficient to confer *in vivo* deacylating activity, as monitored by measuring the APT2-dependent depalmitoylation of ZDHHC6 (**Fig. 6cd**). These observations confirm that the ability of APT2 to deacylate target proteins in cells requires both the ß tongue and acylation of Cys2.

### Identification of a palmitate binding pocket in APTs

To further understand how APTs act on their substrates, we reanalyzed our crystal structures of APT1. We noticed that about one third of the proteins in the asymmetric unit contained a fatty acid in a hydrophobic pocket in the vicinity of the catalytic site (**Fig. 7a, Supplementary Fig.4a**). The electron-density maps specifically identified a palmitic acid (C16:0) with the carboxylate group facing the catalytic Ser119. We therefore extracted the ligand with ethanol and performed ultraperformance liquid chromatography (UPLC) quadrupole-time-of-flight mass spectrometry (Q-TOF-MS) analysis to confirm the presence of the palmitic acid in all three structures (WT, C2S, and S119A APT1) (**Fig. 7b**). Since the fatty acid was not added during expression in *E. coli*, purification, or crystallization, it must have been retained during purification, as was previously observed for other proteins (Ngaki *et al*, 2012).

**Figure 7:**
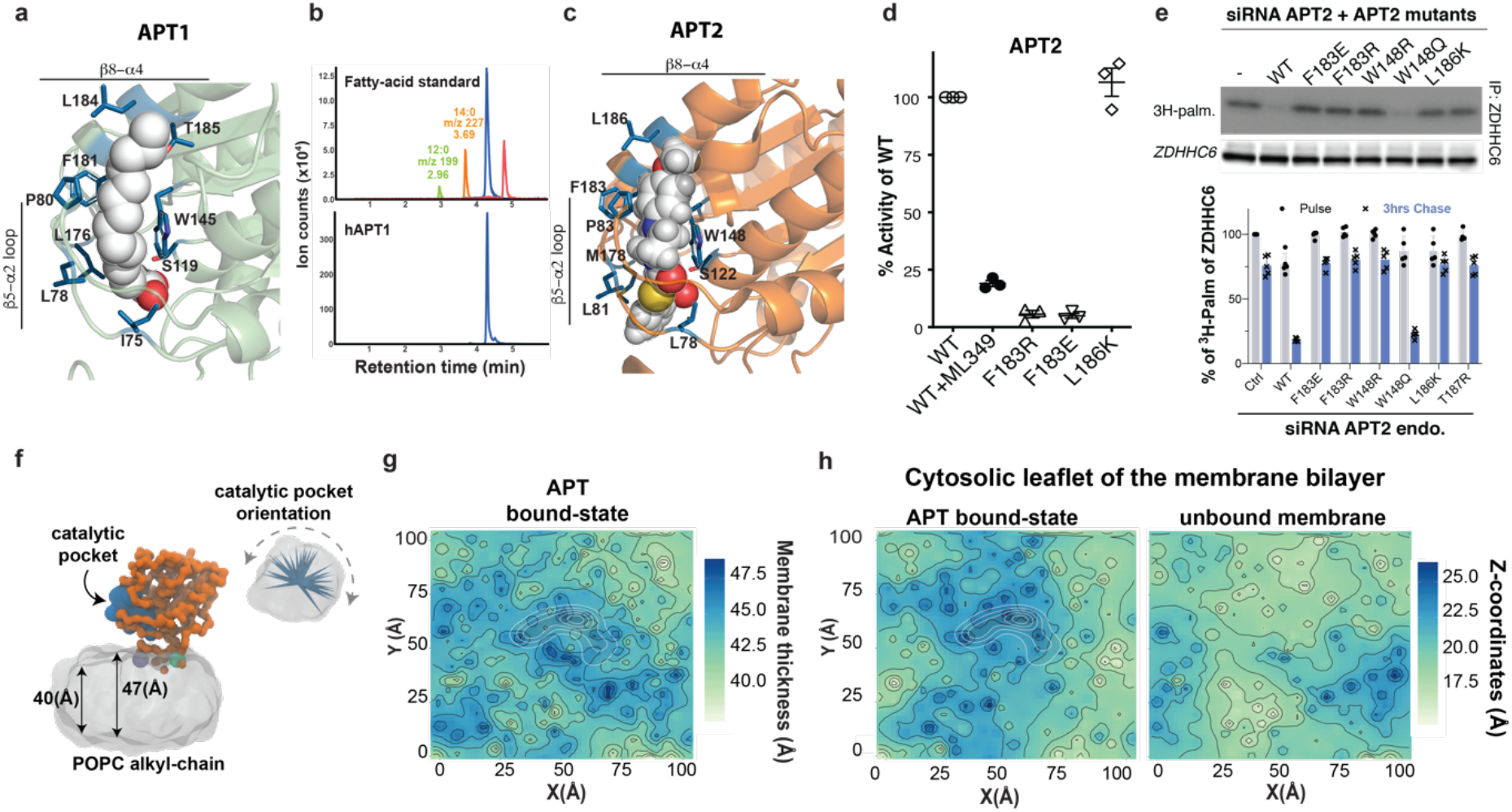
Identification of acyl chain binding pocket in APTs. **a**. Close-up view of the hydrophobic channel of APT1 with palmitic acid. The palmitic acid is represented in van der Waals representation. The side-chain of residues forming the hydrophobic cavity are shown as blue sticks. **b**. Analysis of the ligands associated with purified APT1 enzyme detected by Q-TOF-MS. **c.** Closeup view of the hydrophobic channel of APT2 with ML349. ML349 is represented in van der Waals representation. The side-chain of residues forming the hydrophobic cavity are shown as blue sticks. **d**. The thioesterase activity of the APT2 hydrophobic pocket mutants was determined at 60 min after the addition of substrate and detergent. The APT2-specific inhibitor ML349 was included as a positive control. Technical replicates were averaged within each experiment, and each experiment was then normalized to WT. The average of each experiment was graphed. Two-tailed two-sample unequal variance t-tests were performed on the raw values normalized to the plate. **e**. ZDHHC6 palmitoylation upon overexpression of APT2 pocket mutants. HeLa cells were silenced for 3 days with APT2 RNAi (in 3’ non-coding sequence) and were transfected with plasmids encoding myc-tagged WT ZDHHC6 and the indicated APT2 constructs for 24 h. The cells were then metabolically labeled for 2 h at 37°C with ^3^H-palmitic acid and were chased for 3 h. Proteins were extracted, immunoprecipitated with anti-myc antibodies, separated via SDS-PAGE, immunoprecipitated with anti-myc antibodies, and analyzed by autoradiography (^3^H-palm). The total extracts (40 μg) were immunoblotted with anti-myc antibodies to determine the expression level of WT and mutant APT2. The calculated value of ^3^H-palmitic acid incorporation into ZDHHC6 with WT APT2 was set to 100%, and the mutants were expressed relative to this (n = 3, error bars represent standard deviation). **f**. Representative snapshot of membrane-bound APT2state. In orange, the APT2 protein. The catalytic pocket is shown as blue surface, and the membrane bilayer in grey **g.** MD-averaged membrane-thickness. In white the space explored by the protein during the simulation. **h.** Averaged phosphate Z-coordinates for the APT-bound and APT-unbound states. The dashed line highlighted the membrane bounded region explored by APT during the simulation.

Recently, the structure of the human protein acyltransferase hDHHC20 was solved as an acylated intermediate mimic (Rana *et al*, 2018), which provided insight into the localization of the palmitic acid. The acylated version was obtained by using the covalent ZDHHC inhibitor 2-bromopalmitate (2-BP). Consequently, we wanted to test whether incubating APT1 with 2-BP could be used to fully occupy the hydrophobic pocket in all molecules of the asymmetric unit, to confirm the existence of this acyl chain binding pocket. We first verified whether the catalytic activity of APT1 and APT2 could be inhibited by 2-BP. Remarkably, 2-BP was as efficient in blocking APT2 as its specific inhibitor ML349 (**Supplementary Fig.4b**). Consistently, 2-BP also inhibited the APT2-mediated depalmitoylation of ZDHHC6 (**Supplementary Fig.4cd**). 2-BP also inhibited APT1, but with a lower affinity (**Supplementary Fig.4b**). Nonetheless, the crystallization of APT1 in the presence 2-BP led to the presence of palmitate in all proteins of the unit cell (**Supplementary Fig.1e**).

We next wanted to study the function of this binding site in more detail. The site encompasses a largely non-polar cavity that sequesters the aliphatic chain of the palmitic acid. The residues composing this hydrophobic binding pocket belong to several secondary structural elements. For instance, the entrance of the pocket is located close to the loop between helices S3-S4 (**Supplementary Fig.4f**, Leu78, Ser79, and Pro80) and is solvent accessible. The top of the channel is formed by residues from helix E (Phe181 and Leu184), while the residues Cys144, Trp145, Leu146, and Pro147 contribute to the end of the pocket (**Supplementary Fig.4f**). Finally, several residues near the active site form the rest of the channel: Leu30, Ile75, Ile76, and Leu176 (**Supplementary Fig.4f**). The shape and size of the hydrophobic channel varied between the apo and the palmitate-containing forms of APT1. In the apo form, the hydrophobic channel was somewhat closed, due to the movement of the S3-S4 loop and the adjacent E helix (**Supplementary Fig.4g**). Rearrangement of residues Leu78, Phe181, and Leu184 modulated the surface area of the pocket. The difference between the volumes of the opened (226 Å^3^) and the closed channel conformation (124 Å^3^) corresponds roughly to the rearrangement of the residues responsible for regulating lipid access.

We next modeled palmitate into the corresponding pocket of APT2 (**Fig. 7c**). This allowed us to identify residues involved in the formation of the pocket, such as Trp148, Phe183, Leu186, and The187. We produced recombinant F183R, F183E, and L186K APT2 mutants and tested their deacylation activity *in vitro*. Mutating Phe183 abolished the acylthioesterase activity, while mutating Leu186 had no effect (**Fig. 7d**). For *in vivo* activity testing, we probed a larger panel of mutants. All six mutants were well expressed (**Supplementary Fig.5a**) and underwent S-palmitoylation (**Supplementary Fig.5b**). However, with the exception of the W184Q mutant, all hydrophobic pocket mutants tested (F183E, F183R, W148R, L186R, T187R) failed to depalmitoylate ZDHHC6 (**Fig. 7e** and **S7c**), showing that the hydrophobic pocket of APT2 is essential for its protein acylthioesterase activity.

Altogether, these observations indicate that the acyl chain of a palmitoylated substrate protein needs to access this newly discovered APT hydrophobic pocket for hydrolysis to occur. This means that the acyl chain must first be extracted from the membrane containing the APT2 target protein. We wondered whether APT2 may affect the membrane’s structure in a manner that would favor movement of the acyl chain out of the bilayer. To evaluate this possibility, we monitored the thickness of the lipid bilayer upon interaction of APT2 with the membrane during the MD trajectories (**Fig. 7f**). We observed a correlation between the presence of APT2 and the membrane thickness: while in the absence of APT2, the average membrane thickness is ~40 Å, when APT2 interacts with the membrane, the thickness increased to ~47 Å (**Fig. 7g**). The protein perturbs mostly the upper leaflet of the bilayer by increasing its thickness locally from 17 Å to 25 Å (**Fig. 7h**). Although simulations at this level of theory cannot explore the complete deacylation reaction mechanism, the local membrane deformation promoted by APT2 can facilitate the localization of palmitoylated substrates closer to the active site and favor the extraction of the palmitoylated cysteine residue from the membrane bilayer to be accommodate into the identified hydrophobic pocket.

## DISCUSSION

Though acyl protein thioesterases are critical components of the acylation-deacylation-regulated protein localization and function (Blanc *et al*, 2019; Zaballa & van der Goot, 2018), little is known about how they act in cells and how they are controlled. Combining structural biology, molecular simulations, mutagenesis, and *in vivo* assays, we herein studied the mode of action of the thioesterase APT2. Through this work, we discovered how APT2 approaches membranes to stably associate with them, how it extracts the acyl chain that is to be removed from its targets, and how cells regulate the cellular amounts of APT2 through two counteracting post-translational modifications, S-acylation and ubiquination.

Our structural studies showed that APT2 exposes a mildly hydrophobic loop, the ß tongue, on its surface that allows it to interact with liposomes. Interestingly, the ß tongue is insufficient to mediate stable interactions with membranes in the cellular context, pointing towards some lipid or curvature specificity that remains to be analyzed. We could determine that the stable binding of APT2 to cellular membranes requires two steps: (i) long range electrostatic interactions caused by patches of positively charged residues that allow APT2 to approach membranes and dip in its ß tongue, and (ii) S-acylation of Cys2 by the acyltransferases ZDHHC3 or 7. This S-acylation is required for APT2 to be active in cells, presumably because APT2 needs to have a sufficient dwell time on membranes to encounter its membrane-associated S-acylated target proteins.

The crystallization studies also led us to a hydrophobic pocket in APT2 that can accommodate palmitate. We determined that mutations in this pocket can inhibit the activity of the enzyme, *in vitro* and *in vivo*, without affecting its membrane binding capacity. The importance of this pocket indicates that upon interaction of APT2 with a S-acylated target protein, the target-bound acyl chain transfers from the membrane to the hydrophobic pocket of APT2. This movement positions the thioester bond between the target protein and the acyl chain near the APT2 catalytic site for hydrolysis. An analysis of the MD simulations of the interaction of APT2 with the membrane indicated that beneath the catalytic pocket, APT2 triggers a membrane deformation, pulling up the lipid monolayer towards the protein. Future studies will address whether this APT2-induced membrane deformation is sufficient to trigger acyl chain extraction and capture by the hydrophobic pocket.

The present work also suggests that cells tightly control the levels of cytosolic APT2. APT2 indeed has a build-in mechanism that targets it to degradation when free in the cytosol. This is mediated by the presence of a ubiquitination site (Lys69) within the ß tongue, which is thus inaccessible when the protein is bound to a given organelle. Why excess APT2 is deleterious to the cell remains to be understood.

All together our work reveals the following mode of action for APT2: soluble APT2 weakly interacts with cellular membranes via its ß tongue; in this state, it can detach from the membrane or encounter ZDHHC3 or 7, which can add an acyl chain to Cys2, allowing APT2 to stably bind and explore the membrane in search of potential substrates. Upon encounter with a substrate, APT2 triggers extraction of the acyl chain from the membrane allowing it to move into the APT2 hydrophobic pocket. With this ideal positioning hydrolysis may occur. Membrane bound APT2 can itself undergo deacylation, as suggested by the stabilizing effect of palmostatin B (**Fig. 4c**), and subsequent release from the membrane. APT2 does not have the ability depalmitoylate itself (Kong *et al*, 2013), and thus the involved acyl protein thioesterase remains to be identified. The cycle can then repeat itself on the same or a different membrane compartment. In its cytosolic form, however, APT2 is vulnerable to ubiquitination on Lys69 in the ß tongue, which targets it to the proteasome for degradation.

Overall, these studies on APT2 have shown us how a key regulator of S-acylation functions and is itself controlled. With this information, we can now start addressing the dynamics and spatio-temporal regulation that allow a single enzyme to modify many targets proteins that localize to different compartments of the endomembrane system.

## MATERIALS AND METHODS

### Cell lines

HeLa cells (ATCC) were grown in complete modified Eagle’s medium (MEM, Sigma) at 37°C, supplemented with 10% fetal bovine serum (FBS), 2 mM L-Glutamine, penicillin, and streptomycin.

### Plasmids and transfection

Human influenza hemagglutinin (HA), Myc or CItrin fusions of APT1 or APT2 were inserted into pcDNA3.1-N1. All mutations of APT1 and APT2 were obtained using a Quikchange mutagenesis kit (Stratagene). The Myc fusion of ZDHHC6 was cloned in pcDNA3.1. For the control transfection, we used an empty pcDNA3 plasmid. Plasmids were transfected into HeLa cells for 24 h (2 μg of cDNA/9.6 cm^2^ plate) using Fugene (Promega). For gene silencing, HeLa cells were transfected for 72 h with 100 pmol/9.2 cm^2^ dish of siRNA using the Interferin (Polyplus) transfection reagent. For the control siRNA, we used the following target sequence from the viral glycoprotein VSV-G: 5’-ATTGAACAAACGAAACAAGGA-3’. siRNA against the human APT2 gene were targeted to the 3’ non-coding region (APT2 target sequence: 5’-CAGCTGCTTCTCAGTCATGAA-3’). siRNAs agains all human ZDHHC genes were the same as previously published (Lakkaraju *et al*, 2012).

### Antibodies and drugs

The following primary antibodies are used: mouse anti-Calnexin (Millipore, MAB3126, RRID: AB_2069152), mouse anti-Tubulin (Sigma, T5168, RRID: AB_477579), mouse anti-Actin (Millipore,MAB1501, RRID: AB_2223041), rat anti-HA-HRP (Roche, 12013819001, RRID: AB_390917), mouse anti-Myc (SIGMA, 9E10 M4439, RRID: AB_439694), mouse anti-Ubiquitin (Enzo, P4D1, PW0930, RRID: AB_1181462), rat anti-HA (Roche, 3F10, RRID: AB_390918), rabbit anti-Giantin (Abcam, ab24586, RRID: AB_448163), anti-GM130 (BD, clone35, RRID_398141), anti-FLAG (sigma, M2, RRID_259529). The anti-HA affinity gel (Roche, 1815016001, RRID: AB_390914) or anti-MYC affinity agarose gel (Pierce, PIER20169) were used for immunoprecipitation.

Drugs were used as follows: MG132 at 10 μM (Sigma, C2211), 4 h at 37°C. Palmostatin B at 1 μM (Calbiochem, 178501), indicated time at 37°C. 2-Bromopalmitate (2-BP; Focus Biomolecules, FBM-10-3284) at 100 μM at 37°C during the indicated time.

### Expression and purification of human acyl protein thioesterases (APT) 1 and 2

Human APT1 and APT2 wildtype and mutant proteins were expressed and purified as described elsewhere (Devedjiev *et al*, 2000) with minor modifications. Briefly, BL21 [DE3] (Novagen 70235-3) harboring the appropriate plasmids were grown at 37°C to OD600 = 0.6. Protein expression was induced by the addition of IPTG to 0.8 mM and cultures were grown for an additional 18 h at 16°C. Cells were lysed by sonication in ice-cold H Buffer (50 mM Tris-HCl pH 8.0, 150 mM NaCl), and cellular debris was pelleted by centrifugation at 15,000 × *g* for 30 min at 4°C. The supernatant was loaded on a HisTrap HP (GE LifeSciences 17524802), and protein was eluted in H Buffer containing 250 mM imidazole. Where applicable, the N-terminal His6-tag was removed by the addition of His6-rTEV protease (Life Technologies) and dialyzed in H Buffer for 36 h at 4°C. Cleaved protein was harvested, and DTT and EDTA were added to concentrations of 5 mM and 1 mM, respectively.

### Circular dichroism (CD) spectroscopy

Circular dichroism (CD) spectra of APT2 were collected using a Chirascan V100 (Applied Biophysics) and 1 mm pathlength quartz cuvettes (Hellma 110-QS). Spectra were collected in triplicate at 1 nm intervals between 280 and 207 nm. The temperature of the 6-cuvette holder was monitored and controlled by a Quantum Northwestern CD 250 Peltier system. For denaturation curves, the temperature of the sample was increased from 4°C to 94°C by 2°C intervals; spectra were collected at each temperature following a 30 s equilibration period.

### Thioesterase Activity Assay

The thioesterase activity assay was modified from Kemp et al. (Kemp *et al*, 2013). Recombinant APT was preincubated for 10 min at room temperature with either DMSO or inhibitor in DMSO, where applicable. The reaction was initiated by the addition of 4-methylumbelliferyl palmitate (Santa Cruz Biotechnology sc-214256, PubChem CID 87248) and Pluoronic^®^ F-127 (ThermoFisher P3000MP, PubChem CID24751). Enzyme activity was monitored by the appearance of methylumbelliferone over time (Ex 360 nm, Em 449 nm). The assay was optimized to use 0.4 μM of enzyme, 0.1 mM of substrate, and 0.1% detergent in HBuffer for a 96-well plate format. The following compounds were used as inhibitors: ML348 (TOCRIS 5345, PubChem CID 3238952), ML349 (TOCRIS 5344, PubChem CID 16193817), and 2-bromopalmitate (Fisher Scientific AC218610500, PubChem CID 82145).

### Crystallization and structure determination

Crystals of APT1 WT, S119A, and C2S bound or not to 2-Br-PLM were grown by sitting drop vapor diffusion at 18°C from a 1:1 mixture of protein solution (at ~10 mg/ml in H Buffer with 5 mM of DTT and 1 mM of EDTA) and reservoir solution. For the 2-Br-PLM complex crystals, the protein at 10 mg/ml was incubated with 2-Br-PLM at a final concentration of 5 mM for 10 min at room temperature before setting up the crystallization drops. The respective reservoir solutions contained: 10% w/v of polyethylene glycol (PEG) 8000, 10% w/v PEG 1000, 0.2 M potassium bromide, 01.M sodium acetate at pH 5.5 for WT; 25% of PEG 2000 MME (monomethyl ether), 0.3 M sodium acetate, 0.1 M sodium cacodylate pH 6.5 for C2S and S119A; and 0.1 M imidazole, 0.1 M MES (2-(N-morpholino)ethanesulfonic acid) monohydrate, 20% v/v PEG 500 MME, 10% w/v PEG 2000, 20 mM 1,6-hexanediol, 20 mM 1-butanol, 20 mM 1,2-propanediol, 20 mM 2-propanol, 20 mM 1,4-butanediol, 20 mM 1,3-propanediol, pH 6.5, for C2S bound to 2-Br-PLM. Crystal growth occurred over a period of 5 to 15 days.

Crystals were flash frozen by immersion in liquid nitrogen after soaking in cryoprotectant solution (reservoir solution supplemented with 25% w/v glycerol). X-ray diffraction data were collected on beamline X06DA at the Swiss Light Source (SLS, PSI, Villigen Switzerland) at −173.15°C (100 K). The X-ray wavelengths used were 0.826 Å for WT, 1 Å for C2S and S119A, and 0.919 Å for C2S/2-Br-PLM. Data were indexed, integrated, and scaled with XDS (Kabsch, 2010). Phase determination was carried out by molecular replacement using Phaser (McCoy *et al*, 2007) of CCP4 Suite and the published structure of hAPT1 (PDB: 1FJ2) as a template. Coot (Emsley & Cowtan, 2004) was used for graphical map inspection and manual rebuilding of atomic models. Phenix (Echols *et al*, 2012) was used for structure refinement. The WT crystals contained 6 molecules in the asymmetric unit (AU), while the C2S and S119 mutants pack with 4 and 2 molecules in the AU, respectively. Apo-crystals contained residual palmitic acid molecules originating from *E. coli* in 50% of the molecules in the AU. However, there is full occupancy of the 2-Br-PLM in the C2S/2-Br-PLM complex crystals. The occupancy of 2-Br-PLM was confirmed by Br-anomalous difference maps, as the crystals were radiated with X-rays of the Br-peak energy. Residues T8/P9 to I229/D230 were modelled in all chains except for chain B in the WT crystals, where residues from N5 could be also modeled. There are no Ramachandran outliers. Crystallographic statistics are listed in Supplementary Table 1. Coordinates and structure factors were deposited in the Protein Data Bank (PDB accession codes: 6QGS for wt/hAPT1; 6QGQ for C2S/hAPT1; 6QGO for S119A/hAPT1; and 6QGN for the C2S-2Br-PLM complex crystals).

### Atomistic molecular dynamics simulations

The initial APT1 conformation of the full-length WT APT1 was taken from our crystallographic structures. The resulting model was fully solvated with TIP3P water models (Jorgensen *et al*, 1983) in a water box of dimension 70×70×70 Å^3^ and neutralized by the addition of NaCl at a concentration of 150 mM. The CHARMM36 force field was used for the parametrization of the protein (with CMAP correction). MD simulations were performed with NAMD 2.9 software (Klauda *et al*, 2010). The system was minimized for 1000 steps and equilibrated in the NPT ensemble for 5 ns at 1 atm and 300 K using a time-step of 2 fs. The pCys2/APT1 system was then simulated for 0.3 μs in the NPT ensemble. Snapshots were taken at 0.1 ns time intervals for structural analysis. The system was equilibrated following the protocol reported by Bovigny et al. (Bovigny *et al*, 2015) and was simulated for 0.3 μs in the NPT ensemble. In both simulated systems, the periodic electrostatic interactions were computed using particle Mesh Ewald (PME) summation with grid spacing smaller than 1 Å.

### Coarse-grained molecular dynamics simulations

The atomistic structure pCys2/APT1 and APT1 were coarse grained from our X-ray structures using the Martini script (Bulacu *et al*, 2013) and were initially positioned 60 Å from the membrane using the Insane script (Wassenaar *et al*, 2015), which generates the POPC phosphatidylcholine bilayer and solvent. The pCys parameters were retrieved from a previous CG study (de Jong *et al*, 2013). Five independent replicas, where the protein was aligned with different initial conditions with respect to the membrane, were set up and the resulting systems were minimized and equilibrated for 10 ps in NVT using a timestep of 2 fs. Afterwards, 100 ps of NPT MD with a time step of 20 fs was applied for increasing the temperature and pressure to the range of 300 K and 1 bar, respectively. The temperature was controlled using the Bussi thermostat with a coupling time of 1 ps (Bussi *et al*, 2007), while the pressure was controlled by a weak semi-isotropic coupling with a reference pressure of 1 bar and a compressibility of 3 × 10^-4^ bar^-1^. Production MD trajectories were collected for 4 μs using a time step of 20 fs. All simulations were performed using GROMACS 5.1.2.

### Analysis of fatty-acid binding

Bound ligands were extracted from WT APT1 by the addition of 500 μl of ice-cold HPLC grade ethanol to 100 μl of protein (at 414 μM). The solution was incubated at −20°C for 3 days and was subsequently centrifuged (16,000 x *g* at 4°C) and evaporated under vacuum. The residual material was resuspended in 200 μl of propanol. Chromatographic separations employed an Agilent Zorbax Extended C18 (2.1 mm x 50 mm, 1.8 μm particle size) reversed-phase column run at a flow rate of 0.4 ml/min. A linear gradient was applied with initial and final mobile phase consisting of 50% water:50% acetonitrile and 100% acetonitrile, respectively. The presence of fatty acid was determined by quadrupole-time-of-flight mass spectrometry (Q-TOF-MS) analyses and comparison with a stock solution (Sigma-Aldrich) composed of four different fatty-acid standards (lauric acid [12:0], myristic acid [14:0], palmitic acid [16:0], and stearic acid [18:0]).

### Size-exclusion chromatography with multi-angle laser light scattering

The mass measurements were performed on a Dionex UltiMate3000 HPLC system coupled with a 3-angle miniDAWN TREOS static light scattering detector (Wyatt Technology). Sample volumes of 100 μl of APT1 WT and ATP2 WT at 40 μM were injected into a Superdex 75 10/300 GL column (GE Healthcare) previously equilibrated with 50 mM Tris-HCl pH 8.0, 150 mM NaCl, and 2 mM TCEP at a flow rate of 0.5 ml/min. The data were further analyzed using ASTRA 6.1 software using the absorbance at 280 nm and the theoretical extinction coefficient for concentration measurements.

### Liposome floatation assays

Liposomes composed of L-alpha-phosphatidylcholine (Avanti polar lipids 131 601C), L-alpha-phosphatidylserine (Avanti polar lipids 840032C), and L-alpha-phosphatidylethanolamine (Avanti polar lipids 840026C) were prepared in a 2:2:1 molar ratio. Next, 2 μM of phospholipid mix was suspended in 2 ml of HBS pH 7.4. Then, 50 μg of purified protein were incubated with 200 μl of liposomes or with 200 μl of HBS pH 7.4 (control) for 2 h at 10°C. The protein-lipid suspension was adjusted to 40% sucrose and loaded on the bottom of a sucrose step gradient (40%/30%/10% sucrose). After 1 h of centrifugation at 100 000 × *g*, the interfaces were collected from the top (1: between 10 and 30%; 2: between 30 and 40%; and 3: bottom), and 10 μl of each interface were analyzed by SDS-PAGE and Coomassie blue.

### ^3^H-palmitic acid radiolabeling experiments

HeLa cells were transfected or not with different constructs, incubated for 3 h in Glasgow minimal essential medium (IM; buffered with 10 mM HEPES, pH 7.4) with 200 μCi/ml of ^3^H palmitic acid (9,10-^3^H[N]) (American Radiolabeled Chemicals, Inc.). The cells were washed, and incubated in DMEM complete medium for the indicated time of chase or directly lysed for immunoprecipitation with the indicated antibodies. For immunoprecipitation, cells were washed three times PBS, lysed 30 min at 4°C in the buffer (0.5% Nonidet P-40, 500 mM Tris pH 7.4, 20 mM EDTA, 10 mM NaF, 2 mM benzamidin and protease inhibitor cocktail [Roche]), and centrifuged 3 min at 5000 rpm. Supernatants were subjected to preclearing with G Sepharose beads prior to the immunoprecipitation reaction. Supernatants were incubated overnight with the appropriate antibodies and G Sepharose beads. After immunoprecipitation, washed beads were incubated for 5 min at 90°C in reducing sample buffer prior to 4-20% gradient SDS-PAGE. Gels were incubated 30 min in a fixative solution (25% isopropanol, 65% H2O, 10% acetic acid), followed by a 30 min incubation with the signal enhancer Amplify NAMP100 (GE Healthcare). The radiolabeled products were imaged using a Typhoon phosphoimager and quantified using a Typhoon Imager (ImageQuanTool, GE Healthcare). The images shown for ^3^H-palmitate labeling were obtained using fluorography on film.

### ^35^S-cystein-methionin radiolabeling experiments

HeLa cells were transfected with different Myc-APT2 constructs for 24 h, the cells were starved in DMEM HG devoid of Cys/Met for 30 min at 37°C, pulsed with the same medium supplemented with 140 μCi of ^35^S Cys/Met (American Radiolabeled Chemicals, Inc.) for 20 min, washed, and incubated in DMEM complete medium for the indicated time of chase before immunoprecipitation as for ^3^H-palmitic acid radiolabeling experiments.

### Membrane-cytosol separation

To prepare the post-nuclear supernatant (PNS), HeLa cells were harvested, washed with PBS, and homogenized by passage through a 22 G injection needle in HB (2.9 mM imidazole and 250 mM sucrose, pH 7.4) containing a mini tablet protease inhibitor mixture (Roche). After centrifugation, the supernatant was collected as HeLa PNS.

PNS was centrifuged for 1 h at 100,000 × *g* in a safe-locked Eppendorf tube in a Sorvall MX150 ultracentrifuge. The pellet and supernatant were dissolved in an equal volume of reducing sample buffer and incubated for 5 min at 90°C prior to separation on a 4-20% gradient SDS-PAGE gel.

### Acyl-RAC capture assay

Protein S-palmitoylation was assessed by the Acyl-RAC assay as previously described (Werno & Chamberlain, 2015) with some modifications. HeLa PNS were lysed in buffer (0.5% Triton-X100, 25 mM HEPES, 25 mM NaCl, 1 mM EDTA, pH 7.4, and protease inhibitor cocktail). Then, 200 μl of blocking buffer (100 mM HEPES, 1 mM EDTA, 87.5 mM SDS, and 1.5% [v/v] methyl methanethiosulfonate [MMTS]) was added to the cell lysate and incubated for 4 h at 40°C to block free the SH groups with MMTS. Proteins were acetone precipitated and resuspended in buffer (100 mM HEPES, 1 mM EDTA, 35 mM SDS). For treatment with hydroxylamine (HA) and capture by Thiopropyl Sepharose^®^ beads, 2 M of hydroxylamine was added together with the beads (previously activated for 15 min with water) to a final concentration of 0.5 M of hydroxylamine and 10% (w/v) beads. As a negative control, 2 M Tris was used instead of hydroxylamine. These samples were then incubated overnight at room temperature on a rotating wheel. After washes, the proteins were eluted from the beads by incubation in 40 μl SDS sample buffer with ß mercaptoethanol for 5 min at 95°C. Finally, samples were separated by SDS-PAGE and analyzed by immunoblotting. A fraction of the PNS was saved as the input.

### Immunofluorescence microscopy

Cells were fixed in 4% paraformaldehyde (15 min), quenched with 50 mM NH4Cl (30 min) and permeabilized with 0.1% Triton X-100 (5 min). Antibodies were diluted in PBS containing 1% BSA. Coverslips were incubated for 1 h with primary antibodies, washed three times in PBS, and incubated 45 min with secondary antibodies. Coverslips were mounted onto microscope slides with ProLong^™^ Gold Antifade Mountant (P36930; Invitrogen). Images were collected using a confocal laser-scanning microscope (Zeiss LSM 700) and processed using Fiji^™^ software.

## Supporting information

Upplementary figures

## Acknowledgments

We thank Davide Demurtas and Graham Knott from the BioEM EPFL Core Facility for the EM analysis, Stefania Vossio and Dr. Dimitri Moreau from the ACCESS Geneva screening platform; Laure Menin from EPFL ISIC proteomic facility and the beamline scientists of X06DA at SLS (Villigen Switzerland). We thank all the members of the F.G.v.d.G. lab for their discussions and suggestions. This work was supported by the Swiss National Science Foundation, the Swiss National Centre of Competence in Research (NCCR) Chemical Biology, and the European Research Council under the European Union’s Seventh Framework Programme (FP/2007-2013) / ERC Grant Agreement n. 340260 - PalmERa’.

## Author contributions

Conceptualization, L.A., M. A., P.A.S, M.D.P., G.V.D.G.; Investigation, L.A., M.A., S.H., M.J.M., M.U.A., P.A. S, G. F., F.P.; Funding Acquisition, M.D.P. % G.V.D.G.; Writing–Original Draft M.A., L.A., S. H., M.J.M., M.D.P., G.V.D.G.; Writing–Review & Editing, L.A., M.A., S.H., M.J.M., M.U.A., P.A. S, G. F., F.P., M.D.P., G.V.D.G.; Resources, S.H., L.A.

