## Supplementary material for "Molecular mode of action of an Acyl Protein thioesterase": Upplementary figures

### Supplementary figures

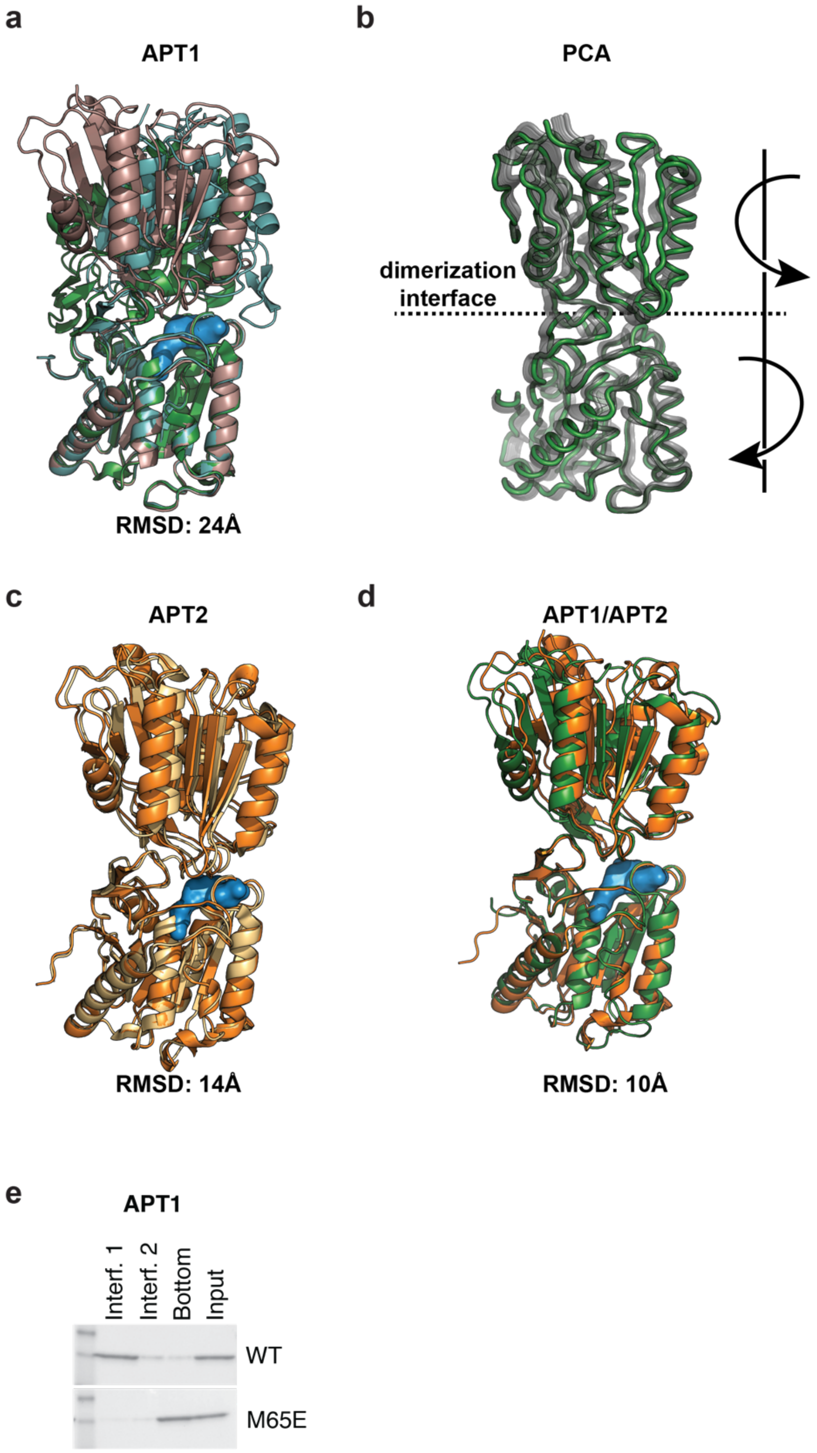

**Supplementary Figure 1**: **The APT crystallographic dimers**. **a**. Comparative superimposition of S119A APT1/6QGO (light-blue), WT APT1/6QGS (green) and WT APT1/1FJ2 (dark-pink). The average backbone RMSD value is 24 Å. **b.** Principal component analysis based on atomistic MD simulations on the WT APT1; **c.** Comparative superimposition of WT APT2/5SYN (orange) and WT APT2/6BJE (light yellow). The average backbone RMSD value is 14 Å; **d.** Superimposition of WT APT1/6QGS (green) and WT APT2/5SYN (Orange). The average backbone RMSD value is 20 Å. In surface blue representation are the catalytic residues: S119, D74 and H208. **e**. Association of WT APT1 and β-tongue mutant with liposomes. WT and mutant APT1 proteins were incubated with liposomes and loaded on the bottom of a sucrose gradient. The different interfaces from the top (1) to bottom were collected, loaded on an SDS-PAGE gel, and revealed with Coomassie blue.

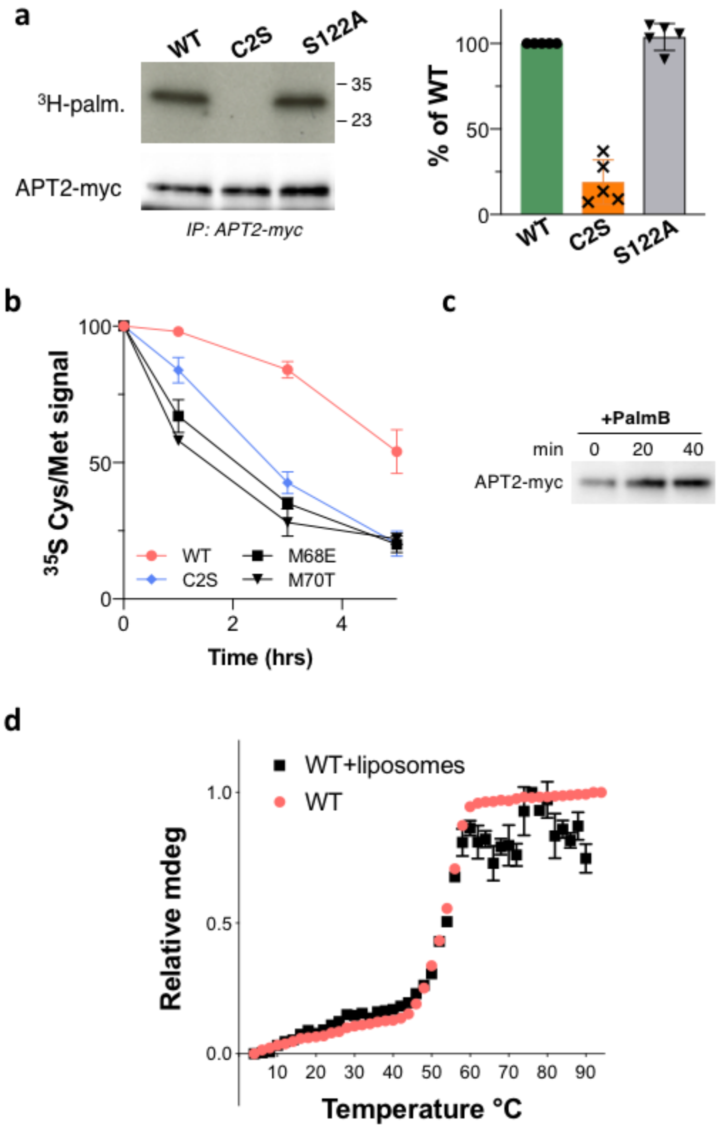

**Supplementary Figure 2: WT and palmitoylation deficient APT2**. HeLa cells were transfected with different myc-tagged APT2 constructs for 24 h. **a**. Cells were then metabolically labeled for 3 h at 37°C with $^{3}$H-palmitic acid. Proteins were extracted, immunoprecipitated with anti-myc antibodies, subjected to SDS-PAGE gel, analyzed by autoradiography ($^{3}$H-palm), and quantified using the Typhoon Imager or by immunoblotting with anti-myc antibodies. The calculated value of $^{3}$H-palmitic acid incorporation into WT APT2 was set to 100%, and the values for the mutants were expressed relative to this (n = 3, error bars represent standard deviation). **b**. Cells were pulsed with $^{35}$S Cys/Met for 20 min and were chased for the indicated time before immunoprecipitation and SDS-PAGE. Degradation kinetics were analyzed by autoradiography, and were quantified using the Typhoon Imager. $^{35}$S-Met/Cys incorporation was quantified for each time point. $^{35}$S-Met/Cys incorporation was set to 100% for t = 0 after the 20 min pulse, and the different chase times were expressed relative to this (n = 3, error bars represent standard deviation). **c**. Cells were incubated with 1 µM of Palmostatin B for different times at 37°C, lysed, subjected to SDS-PAGE, and analyzed by immunoblotting with anti-myc antibodies. **d**. Thermal denaturation profiles of WT APT2 protein alone or associated with liposomes (interface 1 of sucrose gradient) as monitored by circular dichroism at 222 nm. The temperature of the samples was increased from 4°C to 94°C by 2°C intervals. The normalized ellipticity at 222 nm is plotted against temperature.

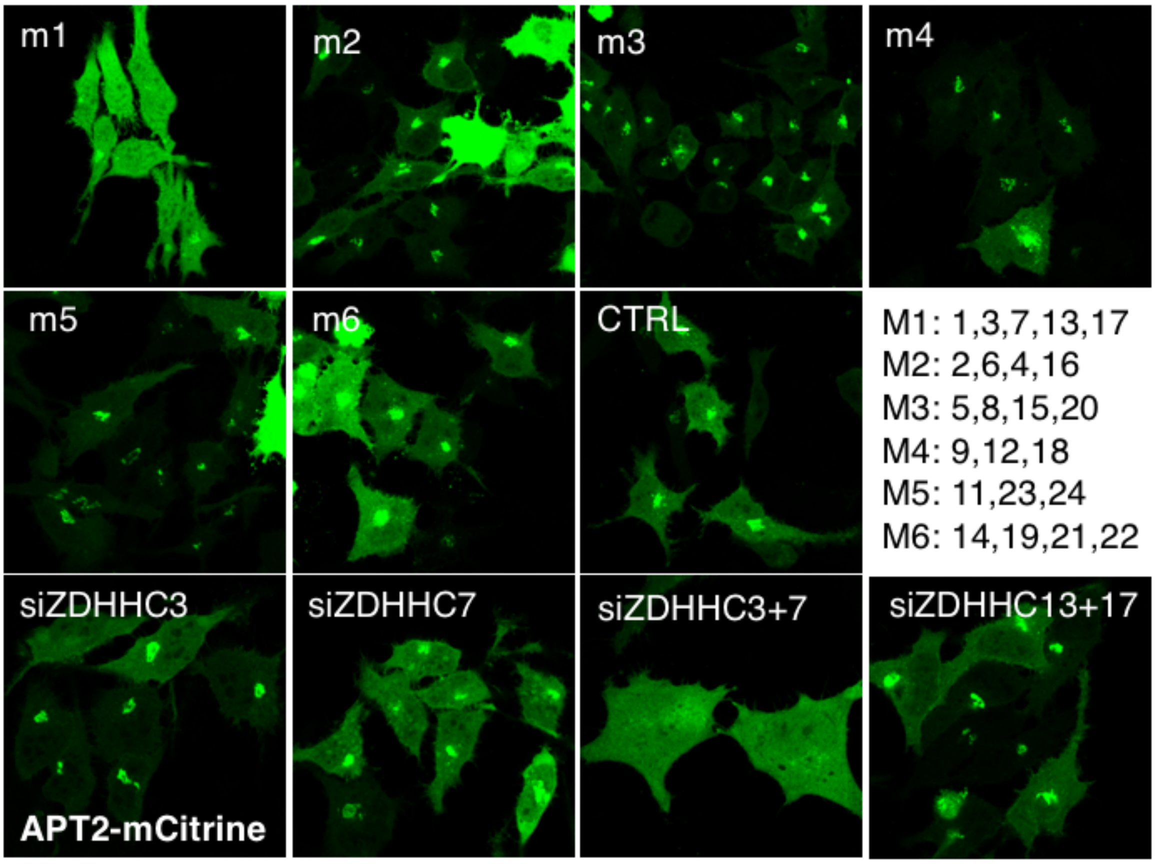

**Supplementary Figure 3: Identification of the APT2 acytransferases.** HeLa cells were silenced for 3 days with individual or mixed pools of ZDHHC RNAi transfected with plasmids encoding citrine-tagged WT APT2. Cells were immunolabeled for citrine-APT2 (green).

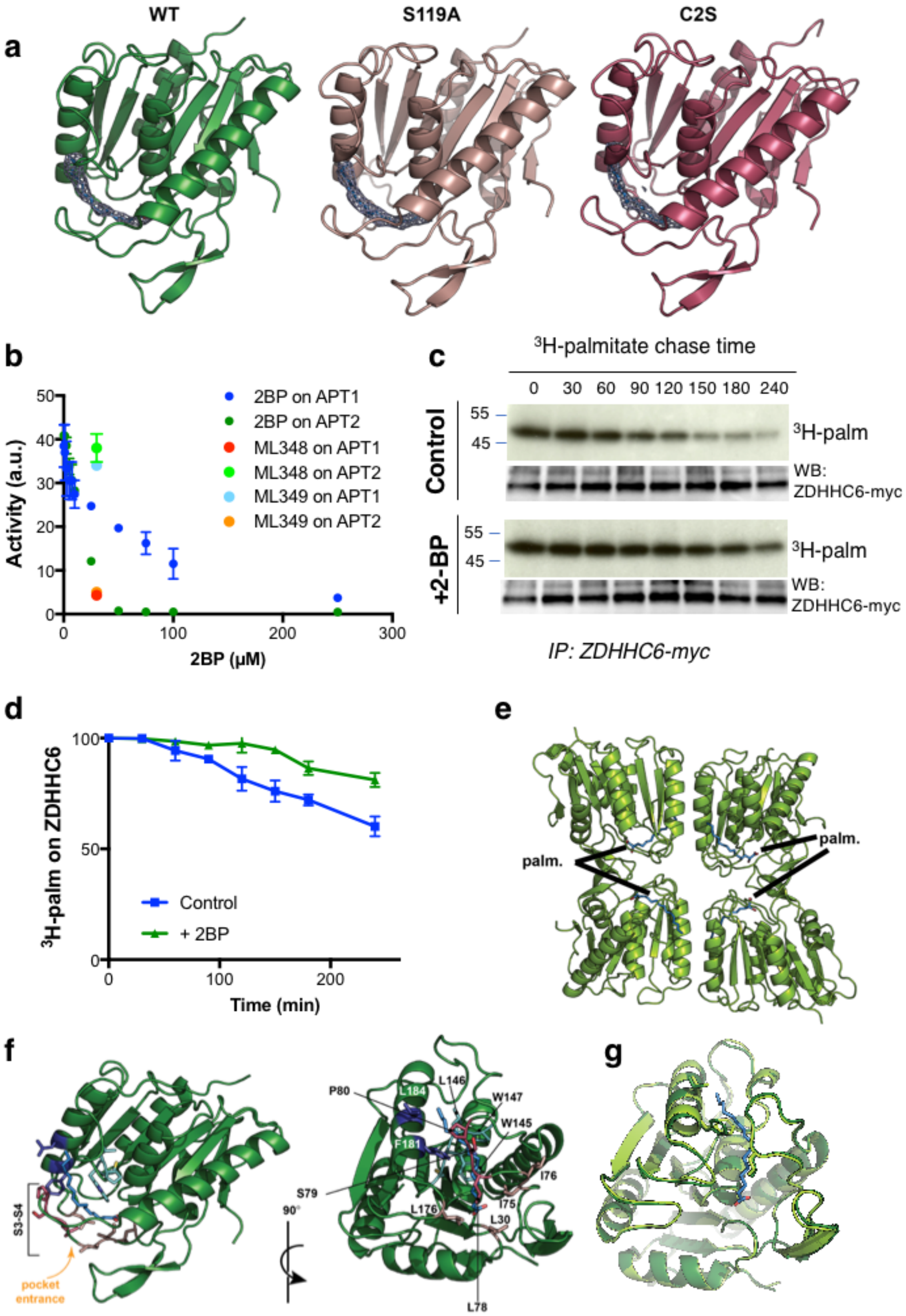

a
WT
S119A
C2S
b
Activity (a.u.)
2BP (µM)
2BP on APT1
2BP on APT2
ML348 on APT1
ML348 on APT2
ML349 on APT1
ML349 on APT2
c
³H-palmitate chase time
0 30 60 90 120 150 180 240
Control
+2-BP
55
45
³H-palm
WB: ZDHHC6-myc
IP: ZDHHC6-myc
d
³H-palm on ZDHHC6
Time (min)
Control
+ 2BP
e
palm.
f
S3-S4
pocket entrance
90°
P80
L184
L146
W147
W145
F181
S79
I76
I75
L176
L30
L78
g

**Supplementary Figure 4**: **Palmitate binding in the APT hydrophobic pocket**. **a**. Ribbon diagrams of APT1 WT and mutants showing the bound palmitate moiety. **b.** Determination of the effect of 2-bromopalmitate on the thioesterase activity of WT APT1 and WT APT2 at 60 min after the addition of substrate and detergent. APT1-specific inhibitor ML348 or APT2-specific inhibitor ML349 at 10 µM were included as positive and negative controls. **cd**. HeLa cells were transfected with plasmids encoding myc-tagged WT ZDHHC6 constructs for 24 h. Cells were metabolically labeled for 2 h at 37°C with $^{3}$H-palmitic acid and were chased for different times in new complete medium in the presence or not of 2-bromopalmitate only during the chase. Proteins were extracted, immunoprecipitated with anti-myc antibodies, subjected to SDS-PAGE, analyzed by autoradiography ($^{3}$H-palm), and quantified using the Typhoon Imager or by immunoblotting with anti-myc antibodies. $^{3}$H-palmitic acid incorporation was set to 100% for cells after the pulse, and values obtained after different chase times were expressed relative to this (n = 3, error bars represent standard deviation). **e**. 2-bromopalmitate (2-BP) was bound in all APT1 subunits in the asymmetric unit. Ribbon diagram of the 2-BP/APT1 asymmetric unit. The 2-BP molecules are shown in blue sticks. **f**. Ribbon diagram of the side (left) and front (right) view of the APT1 enzyme. The residues forming the binding pocket are shown in sticks representation: in red the residues forming the entrance of the pocket, in blue the residues composing the top of the pocket and in light-blue the residues defining the end of the pocket. The rest of the channel is formed by the residues in pink. The entrance of the channel is solvent-exposed and indicated with an orange arrow. The palmitic acid is shown as blue sticks. **g**. Ribbon diagram of the front view of the apo WT APT1 enzyme (light-green) and palmitate-bounded WT APT1 enzyme (dark-green). In sticks representation, the residues involved in the regulation of the lipid access: Leu184, Phe181, and Leu78.

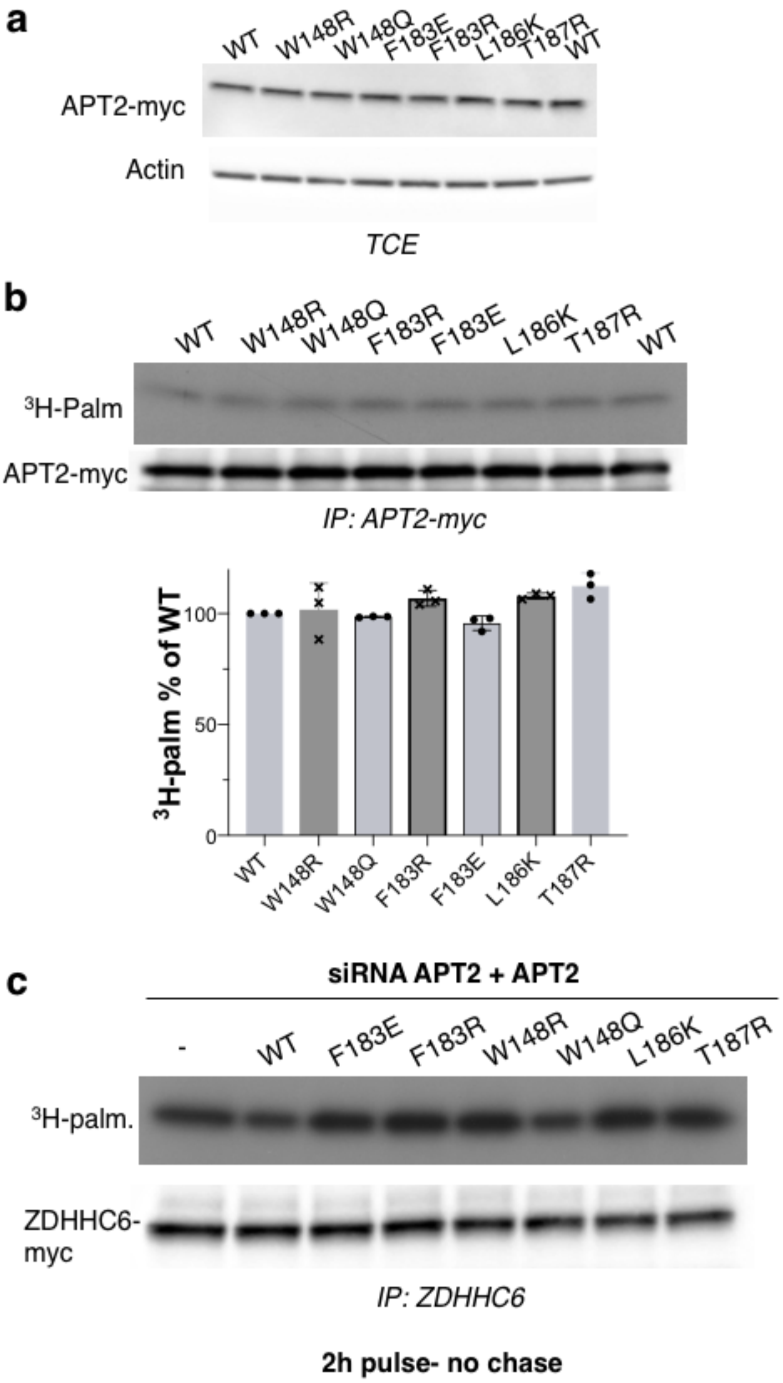

**Supplementary Figure 5**. **APT2 pocket mutants**. **a**. Level of expression of APT2 pocket mutants. Total cell extract (TCE) from cells expressing WT or mutant APT2 for 24 h were subjected to SDS-PAGE, and analyzed by immunoblotting with anti-myc antibodies. Anti-actin antibodies were used as a loading control. **b**. Palmitoylation of APT2 pocket mutants. HeLa cells were transfected 24h with plasmids encoding the indicated APT2 constructs. Cells were then metabolically labeled for 2 h at 37°C with $^{3}$H-palmitic acid. APT2 was immunoprecipitated with myc antibodies, subjected to SDS-PAGE, immunoblotted with anti-myc antibodies, and analyzed by autoradiography ($^{3}$H-palm). Quantification of the $^{3}$H-palmitic acid incorporation into different APT2 mutants. The calculated value of $^{3}$H-palmitic acid incorporation into WT APT2 was set to 100%, and the mutants were expressed relative to this (n = 3, error bars represent standard). **c**. HeLa cells were silenced for 3 days with APT2 RNAi (in 3' non-coding sequence) transfected with plasmids encoding myc-tagged ZDHHC6 and the indicated APT2 constructs for 24 h. Cells were then metabolically labeled for 2 h at 37°C with $^{3}$H-palmitic acid. ZDHHC6 was immunoprecipitated with myc antibodies, subjected to SDS-PAGE, immunoblotted with anti-myc antibodies, and analyzed by autoradiography ($^{3}$H-palm).

**Supplementary Table 1. Data collection and refinement statistics (molecular replacement)**

| | hAPT1 wt | hAPT1 C2S-PLM | hAPT1 S119A | hAPT1 C2S–2BrPLM |
|---|---|---|---|---|
| **Data collection** | | | | |
| Space group | P41212 | P21212 | P64 | P212121 |
| Cell dimensions | | | | |
| *a*, *b*, *c* (Å) | 81.66, 81.66, 441.946 | 146.14, 160.74, 40.68 | 97.89, 97.89, 100.03 | 58.90, 109.61, 174.71 |
| α, β, γ (°) | 90, 90, 90 | 90, 90, 90 | 90. 90. 120 | 90, 90, 90 |
| Resolution (Å)* | 49.95-2.76 (2.92-2.76) | 48.71-2.60 (2.74-2.60) | 48.94-2.60 (2.74-2.60) | 46.43-2.09 (2.21-2.09) |

| | | | | |
|---|---|---|---|---|
| Unique reflections* | 73690 (11724) | 30525 (4352) | 16831 (2432) | 66902 (9483) |
| $R_{meas}$* | 0.13 (0.57) | 0.18 (0.90) | 0.11 (0.73) | 0.10 (0.55) |
| $I / \sigma I$* | 20.1 (5.0) | 9.6 (2.0) | 14.0 (2.4) | 12.8 (3.2) |
| Completeness (%)* | 99.7 (98.1) | 99.9 (99.5) | 99.9 (97.3) | 99.7 (98.3) |
| Redundancy* | 14.0 (13.4) | 7.9 (7.6) | 6.3 (5.9) | 7.7 (7.0) |
| **Refinement** | | | | |
| Resolution (Å) | 49.95-2.76 | 48.71-2.60 | 48.94-2.60 | 46.43-2.09 |
| No. reflections | 73550 | 30461 | 16797 | 66823 |
| $R_{work}$ / $R_{free}$ | 18.4/23.0 | 19.1/23.8 | 18.8/23.6 | 23.0/27.3 |
| No. atoms | | | | |
| Protein | 10007 | 6680 | 3337 | 6731 |
| Ligand/ion | 37 | 53 | 18 | 76 |
| Water | 196 | 132 | 51 | 438 |
| *B*-factors | | | | |
| Protein | 35.6 | 37.4 | 48.4 | 31.5 |
| Ligand/ion | 35.6 | 47.6 | 47.0 | 32.2 |
| Water | 31.5 | 38.0 | 44.6 | 34.4 |
| R.m.s. deviations | | | | |
| Bond lengths (Å) | 0.003 | 0.003 | 0.003 | 0.007 |
| Bond angles (°) | 0.56 | 0.54 | 0.64 | 0.83 |

*Values in parentheses are for highest-resolution shell.
